# Cytosolic Adaptation to Mitochondrial Precursor Overaccumulation Stress Induces Progressive Muscle Wasting

**DOI:** 10.1101/733097

**Authors:** Xiaowen Wang, Frank A. Middleton, Rabi Tawil, Xin Jie Chen

**Affiliations:** Department of Biochemistry and Molecular Biology, State University of New York Upstate Medical University, Syracuse, NY 13210, USA; Department of Neuroscience and Physiology, State University of New York Upstate Medical University, Syracuse, NY 13210, USA; Department of Neurology, University of Rochester, Rochester, NY 14642, USA

## Abstract

Mitochondrial dysfunction causes muscle wasting (or atrophy) in many diseases and probably also during aging. The underlying mechanism is unclear. Accumulating evidence suggests that substantial levels of bioenergetic deficiency and oxidative stress are insufficient by themselves to intrinsically cause muscle wasting, raising the possibility that non-bioenergetic factors may contribute to mitochondria-induced muscle wasting. In this report, we show that chronic adaptation to mitochondria-induced proteostatic stress in the cytosol induces muscle wasting. We generated transgenic mice with unbalanced mitochondrial protein loading and import, by a two-fold increase in the expression of the nuclear-encoded mitochondrial carrier protein, Ant1. We found that the *ANT1*-transgenic mice progressively lose muscle mass. Skeletal muscle is severely atrophic in older mice without affecting the overall lifespan. Mechanistically, Ant1 overloading induces aggresome-like structures and the expression of small heat shock proteins in the cytosol. The data support mitochondrial Precursor Overaccumulation Stress (mPOS), a recently discovered cellular stress mechanism caused by the toxic accumulation of unimported mitochondrial precursors/preproteins. Importantly, the *ANT1*-transgenic muscles have a drastically remodeled transcriptome that appears to be trying to counteract mPOS, by repressing protein synthesis, and by stimulating proteasomal function, autophagy and lysosomal amplification. These anti-mPOS responses collectively reduce protein content, which is known to decrease myofiber size and muscle mass. Our work therefore revealed that a subtle imbalance between mitochondrial protein load and import is sufficient to induce mPOS *in vivo*, and that anti-mPOS adaptation is a robust mechanism of muscle wasting. This finding may help improve the understanding of how mitochondria contribute to muscle wasting. It could have direct implications for several human diseases associated with *ANT1* overexpression, including Facioscapulohumeral Dystrophy (FSHD).

**One Sentence Summary:** Proteostatic adaptations to proteostatic stress in the cytosol caused by unbalanced mitochondrial protein loading and import lead to progressive muscle wasting.

## Introduction

Muscle wasting (or atrophy) is defined by reduction of myofiber size, muscle mass and strength. It occurs during physiological aging, as well as in many diseases including neuromuscular degeneration, cancer, sepsis, diabetes and cardiac failure. The progressive loss of muscle mass with aging is also known as sarcopenia. Sarcopenia affects 35.4% and 75.5% of women and men over 60 years of age, respectively (*1*), accounting for compromised life quality and increased morbidity/mortality in the elderly. Muscle wasting is generally attributed to excessive protein degradation relative to protein synthesis (*2*). Mitochondrial dysfunction is known to cause muscle wasting in many mitochondrial myopathies(*3*). It is also proposed to play a role in the progressive muscle wasting process during aging (*4, 5*). However, how mitochondrial dysfunction contributes to muscle wasting and whether it is mechanistically linked to the loss of protein homeostasis are poorly understood. Interestingly, accumulating evidence indicates that merely reducing mitochondrial respiration and mitochondrial ATP export, along with elevation of oxidative stress, has little or limited effect on muscle mass homeostasis during aging (*6–9*). These interesting findings raise the possibility that mitochondria may affect muscle mass homeostasis by mechanisms additional to or distinct from bioenergetic defect and oxidative stress.

Mitochondria are multifunctional. Mitochondrial abnormalities have been shown to affect cell fitness and survival by disrupting many processes that are not directly related to OXPHOS. These processes include, but are not limited to, protein import, protein quality control, proteostatic signaling, metabolic remodeling, toxic metabolite accumulation, apoptosis, mitochondrial dynamics, phospholipid synthesis, calcium homeostasis, redox balance maintenance, iron-sulfur cluster/heme biosynthesis, cell-non-autonomous signaling and epigenetic regulation (*10–19*). Our recent studies demonstrated that various forms of mitochondrial damage, including protein misfolding and reduced protein quality control, can directly trigger cytosolic proteostatic stress and cell death independent of bioenergetics, a phenomenon named mitochondrial Precursor Over-accumulation Stress (mPOS) (*20*). mPOS is characterized by reduced protein import and the toxic accumulation of unimported mitochondrial precursor/preproteins in the cytosol. Other studies documented cellular responses to a direct impairment of the mitochondrial protein import machinery. It was found that protein import stress in yeast activates different cytosolic mechanisms including the Unfolded Protein Response activated by mistargeting of proteins (UPRam), mitochondrial protein Translocation-Associated Degradation (mitoTAD) and mitochondrial Compromised Protein import Response (mitoCPR), to stimulate proteasomal degradation of unimported proteins and the removal of clogged preproteins on the mitochondrial surface (*21–23*). Expression of genes encoding cytosolic chaperones is activated to counteract the proteostatic stress in the cytosol (*24*). Mitochondrial protein import stress also causes the nuclear and mitochondrial translocation of specific cellular factors to reprogram gene expression and restore cellular homeostasis (*25–27*). Currently, it is yet to be determined as to whether mPOS occurs in animals, how animal cells respond to mPOS *in vivo*, and whether mPOS and anti-mPOS signaling affect the homeostasis and function of animal tissues such as the skeletal muscle.

In cultured human cells, we previously showed that overexpression of mitochondrial carrier proteins is sufficient to unbalance protein load and import, and to induce mPOS and cell death (*28*). In the current report, we show that a two-fold increase in the expression of the mitochondrial carrier protein, Ant1, is sufficient to cause mitochondrial protein overloading relative to import capacity. This results in mPOS. Interestingly, we found that chronic proteostatic adaptation to mPOS drastically remodeled the transcriptome. This unbalances protein synthesis and degradation in the cytosol and ultimately, leads to progressive muscle wasting.

## Results

### Moderate Ant1 overexpression causes progressive muscle wasting

To learn whether mitochondrial protein import is readily saturable *in vivo* and to determine the physiological consequences of protein import stress, we generated transgenic mice overexpressing Ant1, a nuclear-encoded mitochondrial carrier protein primarily involved in ADP/ATP exchange on the mitochondrial inner membrane (IMM). *ANT1*-overexpression is associated with Facioscapulohumeral Dystrophy (FSHD)(*29, 30*). An earlier study reported the use of transgenic mice expressing *ANT1* cDNA from the human α-skeletal actin promoter, as a potential model of FSHD(*31*). These mice did not develop muscle pathology. However, that study failed to examine whether or not the Ant1 protein was actually overexpressed compared to control animals, which was a considerable limitation given that Ant1 is highly abundant in muscle mitochondria(*32*). Thus, whether *ANT1* overexpression does, or does not, cause muscle pathology remains undetermined. To address this, we generated *ANT1*-transgenic mice using the *ANT1* genomic locus and its native promoter (Fig. 1A). We obtained two independent hemizygous transgenic mice (hereafter referred to as *ANT1*^*Tg*^/+). Both were kyphotic (Supplemental Fig. 1A and 1B). We then mainly focused our studies on one of the transgenic lines, in which Ant1 levels are increased by only two-fold in the skeletal muscles (Fig. 1B and 1C). These mice cease to gain body weight at the age of 10-12 weeks, with males being more affected than females (Supplemental Fig. 1C and 1D).

**Fig. 1.**
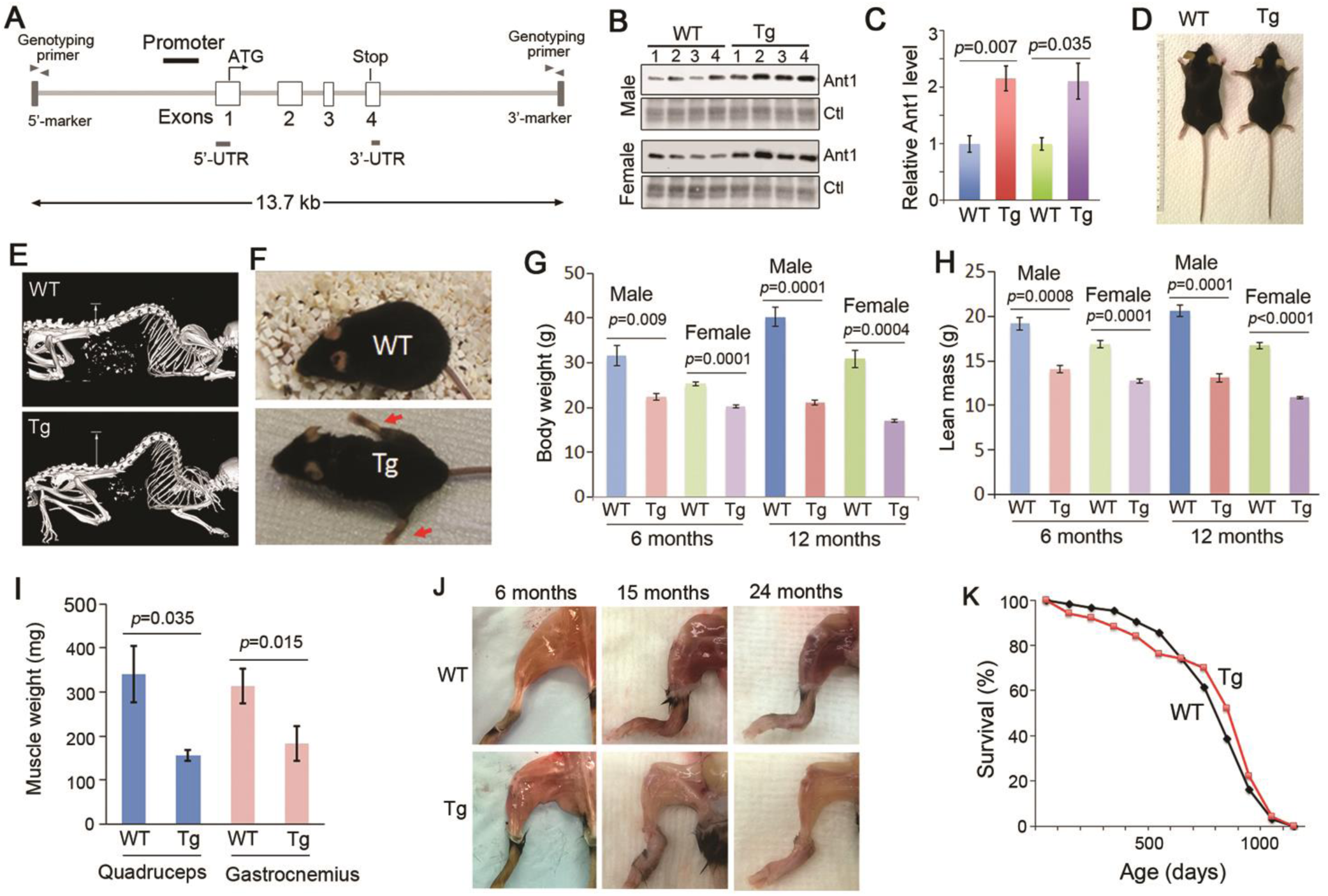
Moderate *ANT1*-overexpression causes progressive muscle wasting without affecting the lifespan. (**A**) Schematic showing the genomic clone of the mouse *ANT1* gene, which was used to generate *ANT1*-transgenic mice. (**B**) and (**C**) Steady state levels of Ant1 in quadriceps muscle from transgenic (Tg) and wild-type (WT) littermates at 6 months old (n=4/genotype/sex), as determined by western blot. Ctl, total protein control. (**D**) Body length at 6 months. (**E**) Micro-CT scanning showing kyphosis at 6 months. (**F**) Gait abnormalities at 19 months. (**G**) and (**H**) Body weight and lean mass at 6 and 12 months (n=4/genotype/sex). (**I**) Loss of quadriceps and gastrocnemius muscles at 14 months. (**J**) Progressive muscle wasting. (**K**) Lifespan of *ANT1*^*Tg*^/+ mice (n=50) compared with wild-type controls (n=62).

We found that the *ANT1*^*Tg*^/+ mice have a normal body length at the age of 6 months (Fig. 1D), suggesting that there is no major developmental delay. Consistent with the kyphotic body morphology, micro-CT confirmed that *ANT1*^*Tg*^/+ mice have increased spinal curvature (Fig. 1E). These mice develop gait abnormalities from the age of 6 months, which progressively deteriorate during aging (Fig. 1F; Supplemental movies 1-3). Parallel to body weight loss (Fig. 1G), quantitative magnetic resonance analysis revealed that the male and female transgenic mice lose lean mass by 26.64% and 24.18% respectively at 6 months, and by 36.28% and 35.01% respectively at 12 months old (Fig. 1H). At 14 months, quadriceps and gastrocnemius muscles are reduced by 54.13% and 41.84%, respectively (Fig. 1I). Skeletal muscle becomes severely atrophic at two years of age (Fig. 1J). Interestingly, the overall lifespan of the *ANT1*^*Tg*^/+ mice is little affected compared with the wild-type controls (Fig. 1K).

To further support the idea that moderate overloading of Ant1 induces muscle wasting, we performed histological analysis of the *ANT1*^*Tg*^/+ and age-matched wild-type mice. We found that the *ANT1*^*Tg*^/+ mice have drastically decreased myofiber size and increased myofiber size variability (Fig. 2A and 2B; Supplemental Fig. 2A and 2B). The average myofiber diameter is reduced by 26.18% and 49.46% at 12 and 20 months old, respectively (Fig. 2C). Round- and angular-shaped myofibers, and mild increase of endomysial connective tissues were observed in the *ANT1*-transgenic muscles (Fig. 2A; Supplemental Fig. 2C-2F). No myofiber type grouping was observed in muscle samples stained for mitochondrial activities, suggesting the lack of chronic neuropathy. Interestingly, we found that the *ANT1*^*Tg*^/+ muscles have increased basophilic stippling, which suggests the induction of possible acidic cellular components (Supplemental Fig. 3A). Moth-eaten myofibers were detected in the *ANT1*^*Tg*^/+ muscles, as indicated by poor staining of NADH, succinate dehydrogenase (SDH) and cytochrome *c* oxidase (COX) (Fig. 2D; Supplemental Fig. 3B). The frequency of myofibers with central nuclei, commonly seen in regenerative myofibers, was not increased in the *ANT1*^*Tg*^/+ muscles compared with controls (Supplemental Fig. 3C), despite that the level of the proapoptotic Bax protein is elevated in response to Ant1 overexpression (Supplemental Fig. 3D and 3E). These data suggest that *ANT1-*overexpression causes progressive muscle wasting mainly by processes that reduce myofiber size.

**Fig. 2.**
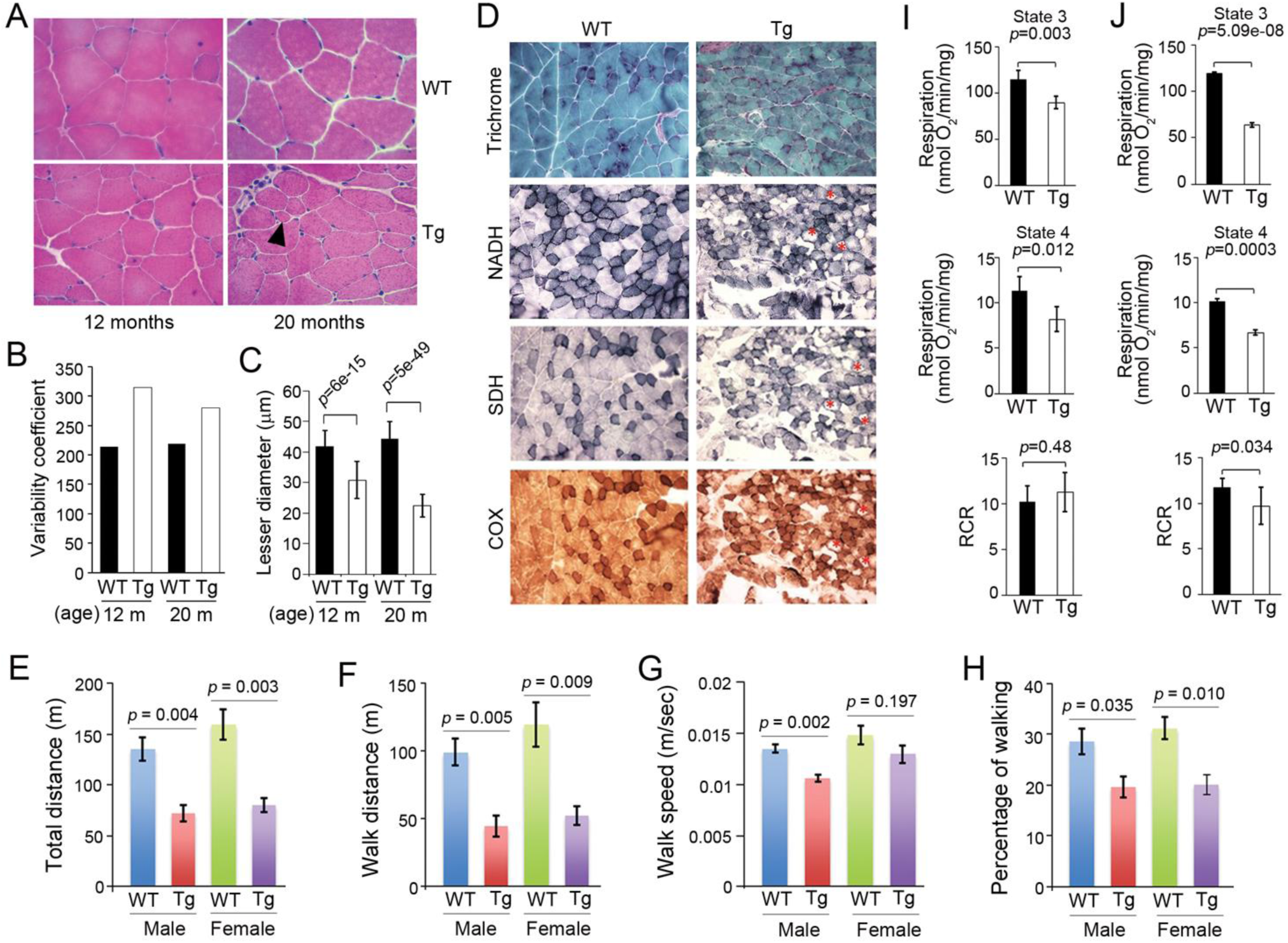
Histological, behavioral and bioenergetic analyses of *ANT1*^*Tg*^/+ mice. (**A**) H&E staining of quadriceps muscles. Arrowhead denotes a rounded small myofiber. (**B**) Variability coefficient of quadriceps myofibers at the age of 12 and 20 months. m, month. (**C**) Lesser diameters of quadriceps myofibers. (**D**) Histology of gastrocnemius muscles from 8.5-month-old *ANT1*^*Tg*^/+ and control mice by modified Gomori Trichrome, NADH, SDH and COX staining. Moth-eaten myofibers are marked by asterisks. (**E-H**) Home cage activity of one-year-old *ANT1*^*Tg*^/+ and littermate control mice in the dark cycle (n=4/genotype/sex). Note that we found no significant differences in home cage activity between *ANT1*^*Tg*^/+ and control mice in the light cycle. Error bars represent S.E.M. and *P* values were calculated by unpaired Student’s *t* test. (**I**) Bioenergetic analysis of skeletal muscle mitochondria from 4-month-old mice. (**J**) Bioenergetic analysis of skeletal muscle mitochondria from 14-month-old mice. Error bars represent standard deviations of four measurements. RCR, respiratory control ratio.

**Fig. 3.**
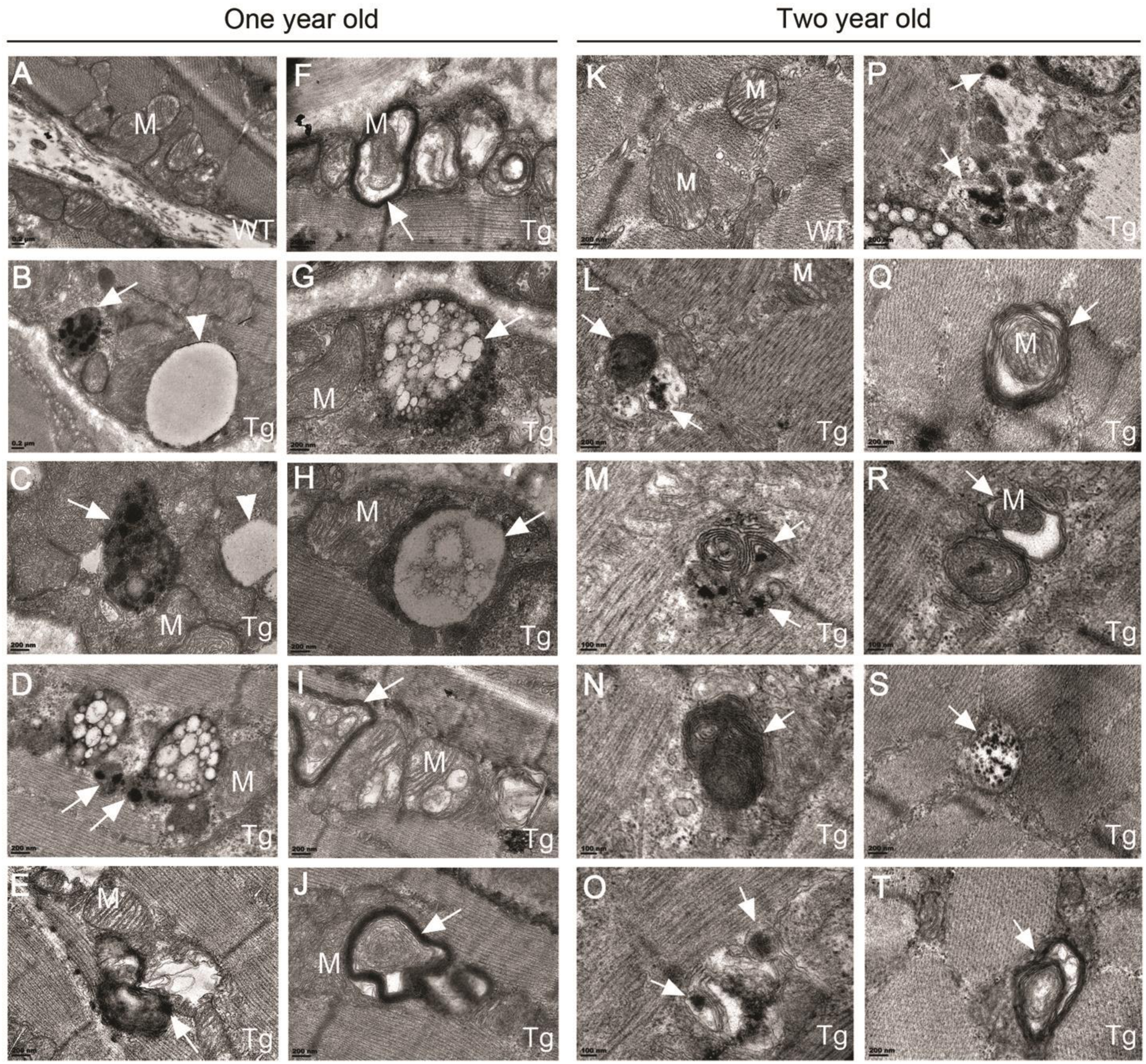
Transmission electron microscopy of quadriceps muscles from *ANT1*^*Tg*^/+ (Tg) and wild-type (WT) mice at one and two years old. M, mitochondria; Arrows denote aggresome/lysosome-like structures that contain electron-dense aggregates (B-E; M-P), mitophagic vacuoles (F, Q and R), multivesicular vacuoles (G and H), multilamellar vesicles (I, J, and T) and structures resembling glycophagy body (S). Arrowheads, structures resembling lipid droplets.

Consistent with muscle wasting and the myopathic phenotypes, the *ANT1*^*Tg*^/+ mice have decreased home cage activities, reflected by reduction in total distance, walk distance, walk speed and the percentage of walking (Fig. 2E – 2H). The effect on walk speed is less pronounced in females compared with males. Using treadmill test, both male and female mice have decreased exercise tolerance (Supplemental Fig. 1E-1H). Taken together, the data indicate that moderate overloading of Ant1 on mitochondria is sufficient to reduce myofiber size, and to cause progressive skeletal muscle wasting and exercise intolerance.

### Mitochondrial respiration is moderately decreased in *ANT1*^*Tg*^/+ muscles

To understand the mechanism by which Ant1 overexpression causes muscle wasting, we first determined mitochondrial respiration in *ANT1*^*Tg*^/+ and control muscles. We found that Ant1 overexpression has a moderate effect on mitochondrial respiration. At 4 months old, the *ANT1* overexpression already has a pronounced effect on body weight (Supplemental Fig. 1C). In the *ANT1*^*Tg*^/+ muscles, the state-3 and state-4 respiratory rates of mitochondria are reduced only by 21.6% and 27.9%, respectively (Fig. 2I). At 14 months, the state-3 and state-4 respiratory rates are reduced by 46.2% and 33.7%, respectively (Fig. 2J). We found that the overall respiratory control ratio (RCR) is unaffected at 4 months old, and is marginally decreased at 14 months old (Fig. 2I and 2J). These data suggest that moderate *ANT1*-overexpression has a relatively mild effect on mitochondrial respiration. Ant1 overloading likely reduces the import of OXPHOS components due to Ant1-induced import stress. A two-fold increase of Ant1 level seems to have very limited effect on respiratory coupling efficiency. Previous studies showed that more severe bioenergetic deficiencies are not sufficient to induce muscle wasting during aging (*6*). We thus speculate that factors other than bioenergetic deficiencies may cause *ANT1*-induced muscle wasting.

### Cytosolic aggresome formation supports mPOS

We recently found that overexpression of mitochondrial carrier proteins in human embryonic kidney HEK293T cells can impose a drastic proteostatic burden in the cytosol, as manifested by the formation of giant aggresomes that contain unimported mitochondrial proteins(*28*). To ascertain whether Ant1 overexpression in *ANT1*^*Tg*^/+ muscles is sufficient to impose protein import stress and induce the formation of cytosolic protein aggregates and/or aggresome structures, we directly examined the *ANT1*^*Tg*^/+ and control muscles by transmission electron microscopy. Indeed, we detected various forms of aggresome-like structures in the cytosol of *ANT1*^*Tg*^/+ but not in that of wild-type (Fig. 3B-3E and 3L-3P). These structures contain a single or multiple electron-dense patches of various sizes and shapes, which reassemble protein aggregates. Many of the aggregates are confined within membrane-encircled vesicles that could derive from the aggrephagic process. These data support the idea that a two-fold increase of Ant1 overexpression is sufficient to saturate the mitochondrial protein import machinery and to cause the accumulation of Ant1, and probably other unimported mitochondrial proteins, in the cytosol.

### Ant1 overloading activates genes involved in mitochondrial protein import and proteostasis, and those encoding small heat shock family B chaperones consistent with mPOS

We speculated that the formation of aggresome-like structures is an adaptive process to counteract the proteostatic stress caused by the overaccumulation of unimported mitochondrial proteins (or mPOS) in the *ANT1*^*Tg*^/+ muscles. To provide additional evidence for mPOS, we analyzed the transcriptome of *ANT1*^*Tg*^/+ skeletal muscles by RNA-Seq. We identified 24,106 genes (Fig. 4A; Supplemental Table 1), among which 2,729 genes were differentially expressed in the *ANT1*^*Tg*^/+ versus wild-type (WT) muscles (one-way ANOVA, *q* < 0.01; Supplemental Table 1). Among these genes, 196 genes were upregulated and 54 genes down-regulated at the ±2.0 fold level, and 42 genes were up-regulated and 6 down-regulated at the ±4 fold level (Fig. 4B; Supplemental Fig. 4). The highly upregulated genes participate in biological functions including Integrated Stress Response (ISR, see below), amino acid transport and metabolism (e.g., *SLC7A1*, *SLC7A5* and *ASNS*), one-carbon metabolism (e.g., *MTHFD2* and *PSAT1*) and myokine signaling (e.g., *FGF21* and *GDF15*). Activation of these genes has previously been reported in various cell and mouse models of mitochondrial stresses and diseases (*33–38*). Gene set enrichment analysis (GSEA) revealed that expression of genes involved in various proteostatic processes in the cytosol is altered (Supplemental Table 3). Many of these genes participate in mRNA processing, ribosomal biogenesis, protein translation and amino acid metabolism. Consistent with our evidence of a mild bioenergetic defect, genes directly involved in oxidative phosphorylation and anti-oxidant defense were not over-represented among the upregulated genes in the *ANT1*^*Tg*^/+ muscles. Instead, we found that several upregulated genes in the “mitochondrial inner membrane” gene ontology group are involved in mitochondrial protein import, including *TOMM40, TOMM5, TOMM34, TIMM8A1, TIMM10, TIMM44, HSPA9, DNAJA3* and *CHCHD10* (Fig. 4C). This supports the existence of a retrograde regulatory mechanism in the skeletal muscle that promotes mitochondrial protein import. *HSPE1*, *LONP1* and *CLPP*, encoding the mitochondrial Hsp10, the Lon protease and the proteolytic subunit of the Clp peptidase in the matrix respectively, were also upregulated (Fig. 4C). This is reminiscent of the mitochondrial Unfolded Protein Response (mtUPR) mechanism that is activated to cope with increased proteostatic stress inside mitochondria (*22, 26*).

**Fig. 4.**
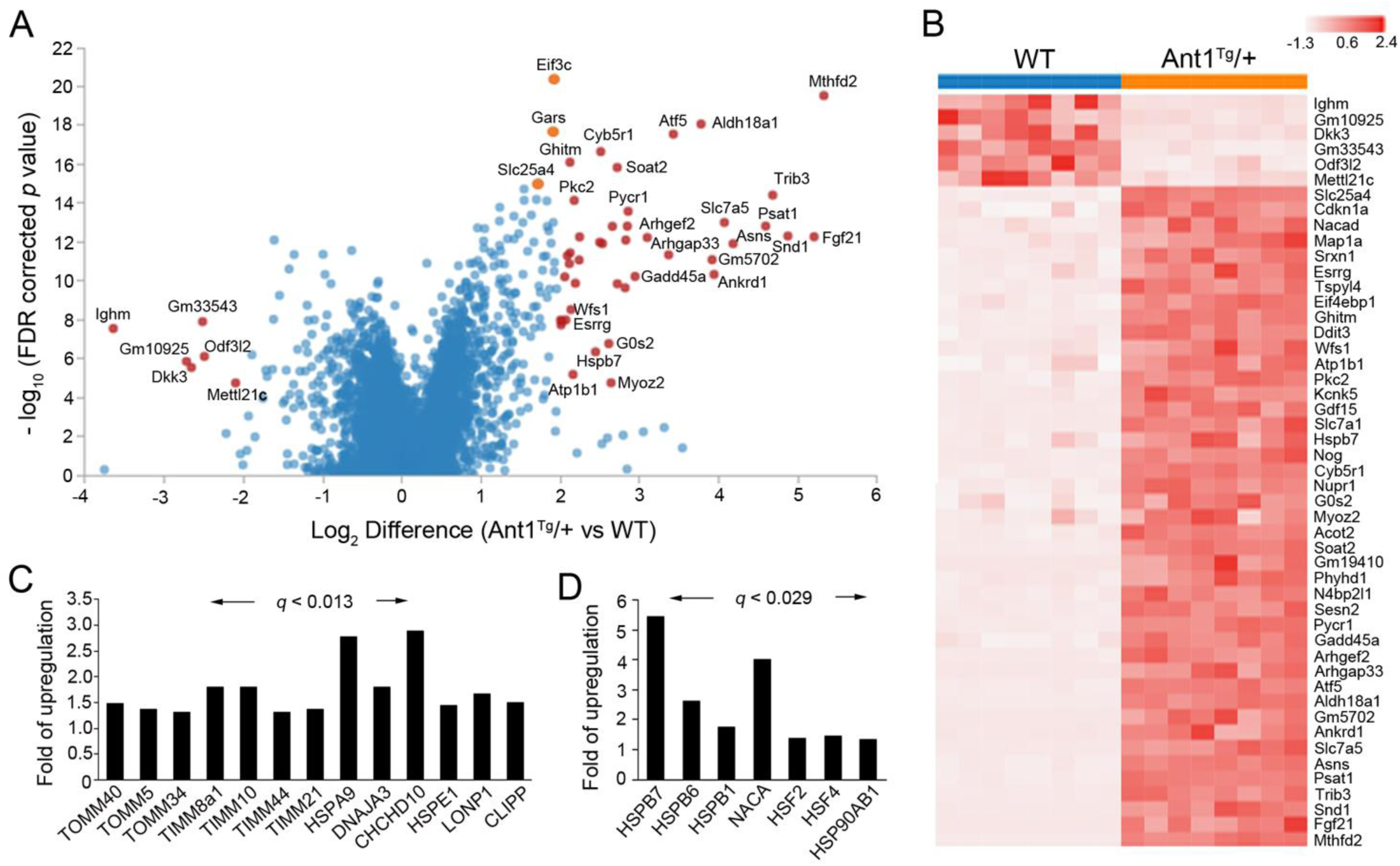
Transcriptional responses to *ANT1*-overexpression determined by RNA-Seq (n=4/genotype/sex) (**A)** Volcano plot with limma-based *p* values. Genes with 4-fold changes at FDR < 0.01 are indicated in red. Additional genes of interest are noted in orange. RNA samples were extracted from quadriceps muscle at the age of 6 months (one-way ANOVA, *q* < 0.01). (**B**) Hierarchical clustering heatmap of 48 genes with 4-fold changes in Ant1^Tg^/+ versus WT mice, at FDR < 0.01. *ANT1* (Slc25a4) is included for comparative purposes. (**C**) Upregulation of genes involved in mitochondrial protein import and proteostasis. (**D**) Upregulation of genes encoding cytosolic chaperones.

In addition to the activation of genes involved in mitochondrial protein import and proteostasis, we found that genes encoding specific cytosolic chaperones and the heat shock transcription factor 2 and 4 (Hsf2 and Hsf4) are upregulated in the *ANT1*^*Tg*^/+ muscles (Fig. 4D). These genes notably encode the small heat shock family B chaperones such as HspB7, HspB6 and HspB1. They are known to provide protection against muscular cell dysfunction and atrophy, and are involved in many neurological, muscular and age-related diseases in humans (*39*). Functionally, the small cytosolic chaperones constitute the first line of cellular defense against proteostatic stress through their activities in capturing early-unfolding states of proteins, and promoting the formation of aggregates to alleviate cytotoxicity (*40*). Finally, we found that *NACA* and *HSP90AB1* are activated in *ANT1*^*Tg*^/+ muscles. *NACA* encodes a subunits of the nascent polypeptide-associated complex (NAC) that plays a role of chaperone under proteostatic stress (*41*). *HSP90AB1* encodes a member of the Hsp90 family chaperones. Taken together, our data support that *ANT1*-overexpression causes protein import stress on mitochondria, which subsequently induces mPOS in the cytosol. We speculate that the small heat shock proteins and other chaperones may play a role in the deposition of unimported mitochondrial proteins.

### Transcriptomic remodeling to suppress protein synthesis in *ANT1*^*Tg*^/+ muscles

Gene set enrichment analysis (GSEA) of the RNA-Seq data revealed changes to expression of genes in several biosynthetic pathways (Supplemental Table 3). Particularly, genes involved in mRNA processing, amino acid biosynthesis, aminoacyl-tRNA biosynthesis and ribosomal biogenesis are significantly over-represented in the *ANT1*^*Tg*^/+ muscles. These changes are typical signatures of amino acid starvation response, possibly reflecting a protein depletion stress in the muscle. Consistent with this, several lines of evidence suggest that the transcriptome of the *ANT1*^*Tg*^/+ muscles is remodeled to suppress protein translation, likely as an adaptive response to counteract mPOS. First, the transcription of *ATF4, ATF5, DDIT3* and *NURP1* is strongly activated (Fig. 5A). These genes are known to be activated as an important regulatory circuit in the Integrated Stress Response (ISR), an elaborating signaling network that is stimulated by divers cellular stresses to decrease global protein synthesis and to activate selected genes in the benefit of cellular recovery (*42*). Supporting this, the phosphorylation of eIF2α, the central player in the ISR, was increased in *ANT1*^*Tg*^/+ muscles (Fig. 5B and 5C). This eIF2α phosphorylation likely upregulates *ATF4*, *ATF5*, *DDIT3* and *NUPR1* (Fig. 5A), by promoting cap-independent translation of these transcriptional factors followed by a feed-forward loop of transcriptional activation(*42*). Activation of these transcriptional factors has been observed in various models of mitochondrial stress (*14, 43–46*). More importantly, phosphorylated eIF2α is known to directly suppress global protein synthesis by interfering with cap-dependent translation initiation, in response to various stresses including mitochondrial damage (*47, 48*).

**Fig. 5.**
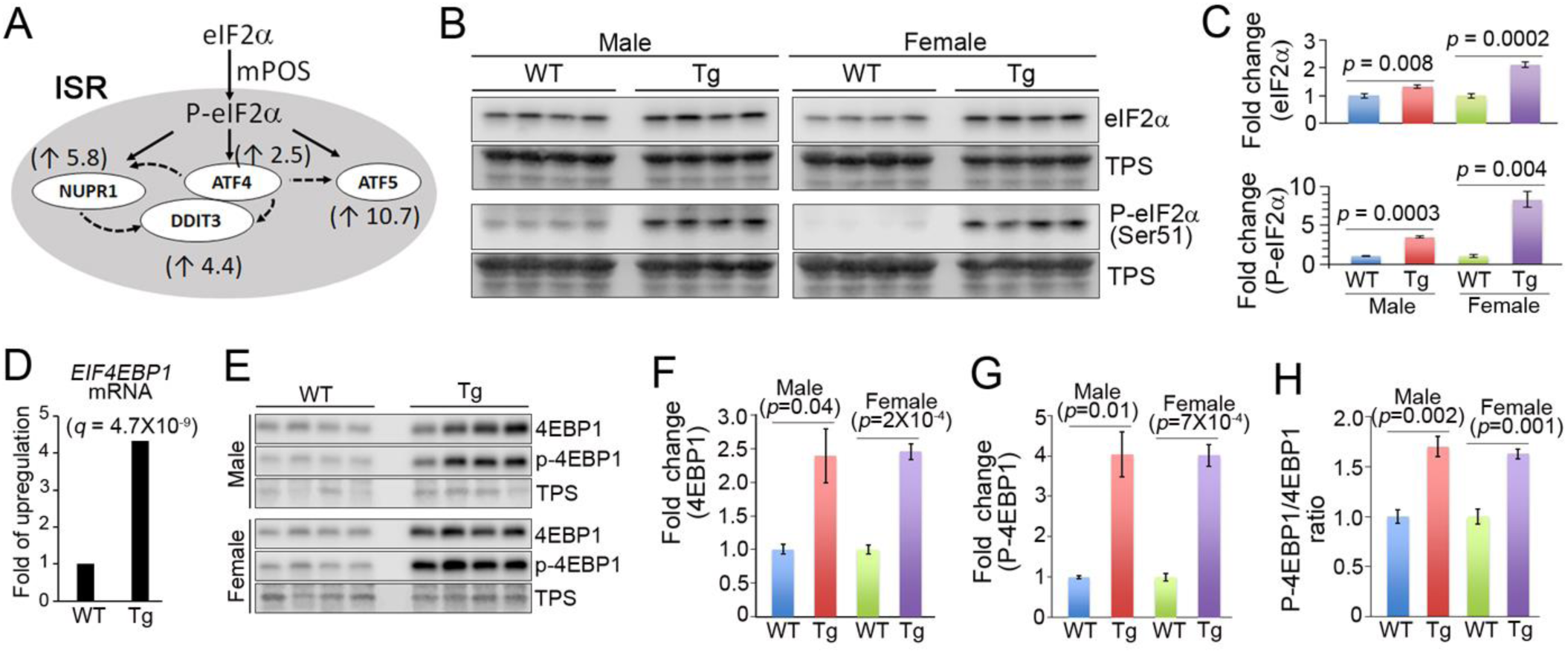
Activation of integrated stress response (ISR) and transcriptional upregulation of *4EBP1* in *ANT1*^*Tg*^/+ (Tg) muscles compared with-wild type (WT) controls (n=4/genotype/sex) **(A)** Upregulation of ISR genes in *ANT1*^*Tg*^/+ muscles. Fold changes in transcription are indicated in the parentheses. Solid lines represent translational activation and dashed lines denote transcriptional activation. (**B**) and (**C**) Western blot showing the levels of eIF2α and its phosphorylated form, P-eIF2α (Ser51), in *ANT1*^*Tg*^/+ and wild-type mice at 6 months old. TPS, total protein staining. (**D**) Transcriptional activation of *EIF4EBP1* in *ANT1*^*Tg*^/+ muscles as determined by RNA-Seq. (**E** – **G**) Western-blot showing the upregulation of 4EBP1 protein and its phosphorylated form, P-4EBP1. (**H**) The ratio between phosphorylated and non-phosphorylated forms of 4EBP1 in *ANT1*^*Tg*^/+ muscles compared with wild-type controls.

Secondly, we found that the transcription of *EIF4EBP1* (or *4EBP1*) is drastically increased in *ANT1*^*Tg*^/+ muscles (Fig. 5D). Consequently, the levels of both 4EBP1 and its phosphorylated form, P-4EBP1, are increased (Fig. 5E-5G). Only the non-phosphorylated 4EBP1 actively binds eIF4E to represses protein translation. Therefore, the transcriptional activation of *4EBP1* would be expected to repress protein translation in the *ANT1*^*Tg*^/+ muscle, even though the P-4EBP1/4EBP1 ratio is moderately increased in the transgenic mice (Fig. 5H). It is noteworthy that 4EBP1 is a substrate of several protein kinases including the mechanistic Target of Rapamycin (mTOR) kinase. We found that the phosphorylation of Rps6 kinase (S6K), another substrate of mTOR, is not significantly increased in the *ANT1*^*Tg*^/+ muscles (Supplemental Fig. 5A – 5E). The data do not support a global increase of mTOR-signaling.

### Activation of multiple protein degradation processes and reduced protein content in *ANT1*^*Tg*^/+ muscles

Our RNA-Seq analysis also revealed that several proteolytic processes are activated in *ANT1*^*Tg*^/+ muscles. First, we found that the transcription of genes encoding proteasomal subunits, *NFE2L1* and *NFE2L2* are upregulated (Fig. 6A & 6B). *NFE2L1* and *NFE2L2* activate the transcription of proteasomal genes. Accordingly, proteasome-associated trypsin-like activity is increased in the *ANT1*^*Tg*^/+ muscles (Fig. 6C). We found that the overall levels of ubiquitinated proteins, and p62 that promotes the formation and removal of ubiquitinated and aggregated proteins, are not increased in the *ANT1*^*Tg*^/+ muscles (Supplemental Fig. 5F-5I). It is likely that activation of proteasomal function, perhaps together with other proteolytic processes (see below), can efficiently prevent the accumulation of ubiquitinated proteins. In acute models of muscle atrophy, *FBXO32* and *TRIM63*, encoding the MAFBx/atrogin-1 and MuRF1 ubiquitin ligases respectively, are frequently upregulated (*49*), Interestingly, we found that these genes are instead down-regulated in the *ANT1*^*Tg*^/+ muscles (Fig. 6D). This observation suggests that muscle wasting in *ANT1*^*Tg*^/+ mice is independent of *FBXO32* and *TRIM63*.

**Fig. 6.**
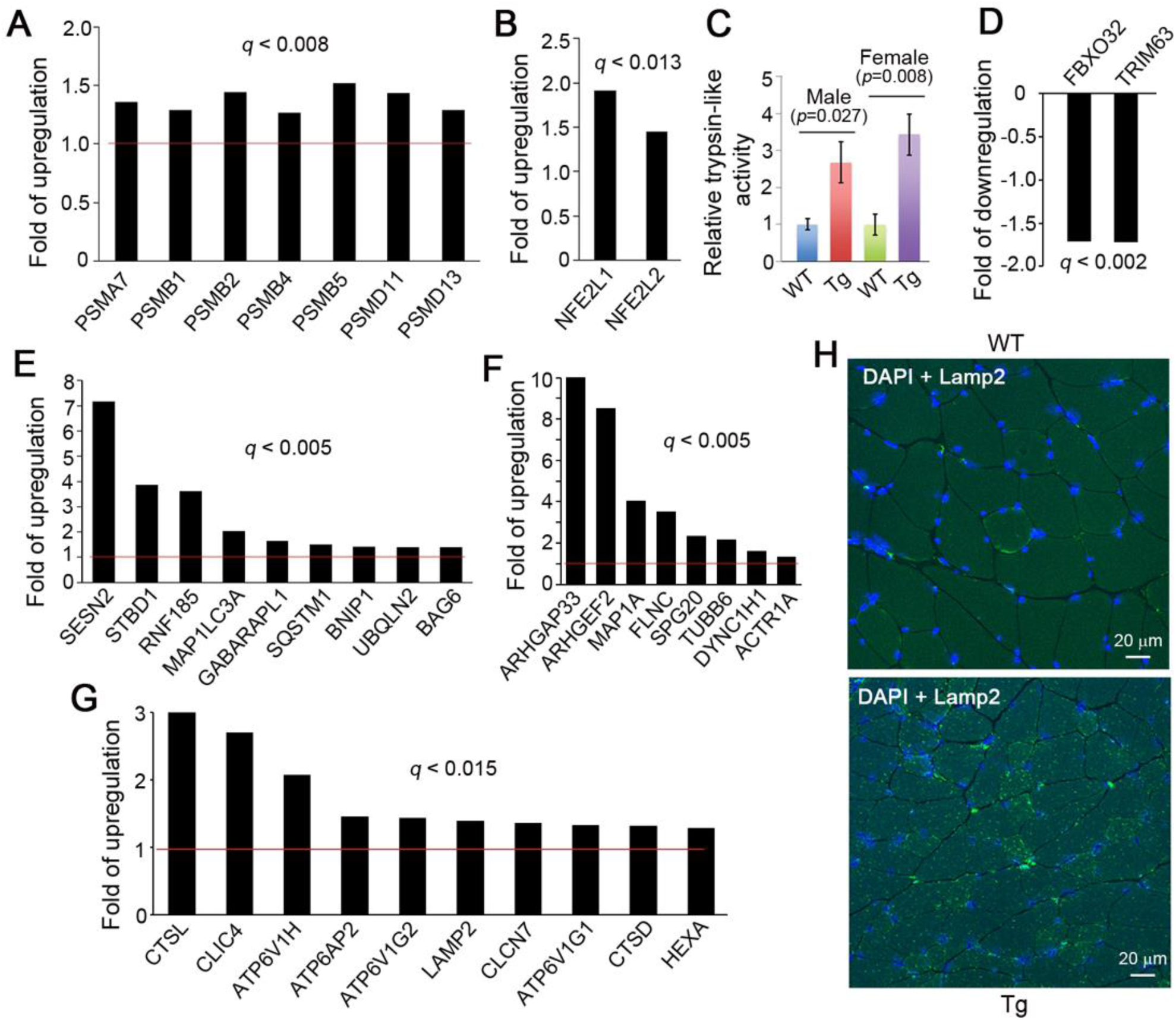
Activation of cytosolic proteolytic pathways in *ANT1*^*Tg*^/+ muscles. (**A**) Moderate transcriptional upregulation of genes encoding proteasomal subunits at 6 months old. (**B**) Transcriptional upregulation of *NFE2L2* and *NFE2L1*. (**C**) Trypsin-like activity associated with the proteasome in *ANT1*^*Tg*^/+ quadriceps muscles. Error bars represent S.E.M. of four male and female mice with four measurements from each mouse. The *p* values were calculated by unpaired Student’s *t* test. (**D**) Transcriptional down-regulation of *FBXO32* and *TRIM63*. (**E**) Transcriptional upregulation of genes involved in autophagy. (**F**) Transcriptional upregulation of genes involved in cytoskeletal organization and intracellular trafficking. (**G**) Transcriptional upregulation of genes encoding lysosomal function. (**H**) Immunofluorescence detection of Lamp2-positive structures in cross-sections of quadriceps from *ANT1*-transgenic and control mice. Blue, DAPI-stained nuclei; green, Lamp2-positive structures.

Secondly, we found that numerous genes involved in autophagy, cytoskeletal organization and intracellular trafficking are upregulated in *ANT1*^*Tg*^/+ muscles (Fig. 6E and 6F). Notably, *SESN2* has been shown to activate mitophagy (*50, 51*). In accordance with this, we detected mitophagic structures in *ANT1*^*Tg*^/+ but not control muscles (Fig. 3F, 3Q and 3R). *STBD1* encodes a selective autophagy receptor for glycogen (*52*), consistent with the presence of glycophagy-like structures in the *ANT1*^*Tg*^/+ muscles (Fig. 3S). *ARHGAP33* and *ARHGEF2* encode a sorting nexin family member and a Rho-Rac guanine nucleotide exchange factor respectively, which are involved in vesicular trafficking (*53–55*). Our transmission electron microscopy analysis also revealed the presence of many vesicular structures of various morphologies in *ANT1*^*Tg*^/+ muscles, which includes multivesicular vacuoles (Fig. 3G and 3H) and multilamellar vesicles (Fig. 3I, 3J and 3T). The origin of these membranous structures remains unclear.

Finally, we found that a group of genes encoding lysosomal function are upregulated in *ANT1*^*Tg*^/+ muscles (Fig. 6G). This includes *CTSL*, encoding the lysosomal cathepsin L protease which plays a major role in the terminal degradation of proteins that are delivered to lysosomes by processes including autophagy. Using immunofluorescence staining, we found that Lamp2-possitive lysosomes and/or lysosome-derived structures are amplified in the *ANT1*^*Tg*^/+ muscles (Fig. 6H; Supplemental Fig. 6; Supplemental movie 4 and 5).

Given the upregulation of multiple proteolytic pathways, together with the activation of ISR and *4EBP1* that repress global protein translation, we speculated that *ANT1*^*Tg*^/+ muscles may have unbalanced protein synthesis versus degradation. This turned out to be the case. We found that the protein content is drastically reduced in the *ANT1*^*Tg*^/+ muscles compared with wild-type controls (Fig. 7A), providing a mechanistic explanation for the muscle wasting phenotype.

**Fig. 7.**
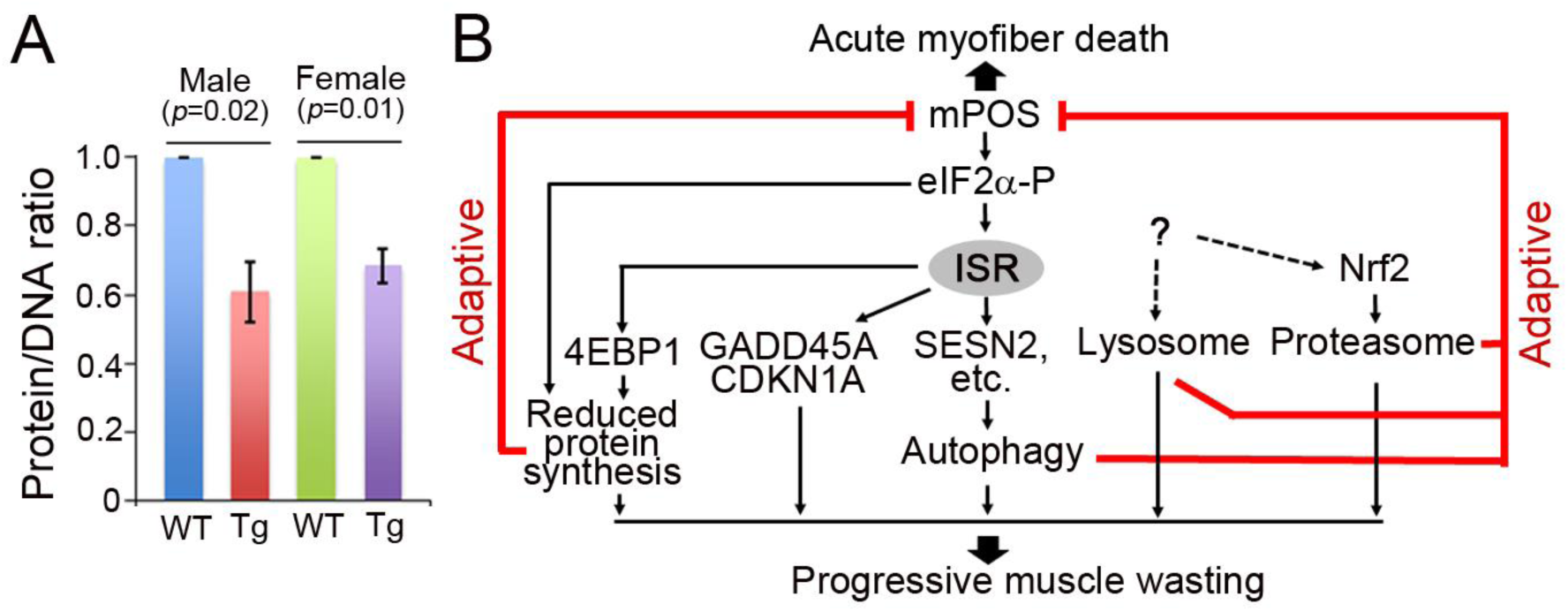
Reduced protein content in *ANT1*^*Tg*^/+ muscles and model for mPOS-induced muscle wasting. (**A**) Protein/DNA ratio in six-month-old quadriceps muscles (n=4/genotype/sex). (**B**) A model for muscle wasting. mPOS-induced proteostatic adaptations unbalance protein synthesis and degradation, which ultimately causes progressive muscle wasting as an undesired trade-off. Red lines represent adaptive processes to counteract mPOS and the dashed lines denote an unknown transcriptional activation mechanism.

## Discussion

Muscle wasting occurs under many disease conditions and during aging. A prevailing mechanism involves an imbalance between protein synthesis and degradation. This reduces protein content, and causes myofiber shrinkage and muscle mass loss (*49*). Mitochondrial dysfunction is known to cause muscle wasting in mitochondrial myopathic diseases (*3*). It is also proposed to contribute to sarcopenia (*4, 5*). However, how mitochondrial dysfunction triggers the atrophying process and whether it involves proteostatic imbalance remain unanswered. In this report, we established a novel mouse model of mitochondria-induced progressive muscle wasting, by moderately overexpressing the Ant1 protein which induces mitochondrial protein import stress. As the lifespan of the animal is unaffected, the scope and mechanism of muscle wasting can be followed in detail during aging. We found that *ANT1* overexpression causes a drastic muscle wasting phenotype in aged mice. Mechanistically, *ANT1* overexpression induces cytosolic proteostatic stress, or mPOS, as supported by aggresome formation and the activation of specific protein chaperones in the cytosol. Muscle cells respond to mPOS by activating multiple proteostatic pathways. First, eIF2α phosphorylation is increased, which is expected to directly repress cap-dependent protein translation. Secondly, we observed the transcriptional activation of 4EBP1, a target of the ISR network (*56*). This leads to the accumulation of both non-phosphorylated and phosphorylated forms of the protein, with the former acting as a repressor of protein translation. Thirdly, we found that Ant1-induced mPOS activates many genes involved in autophagy, vesicular trafficking, lysosomal biogenesis and proteasomal function. Activation of these genes may be protective against mPOS, by directly contributing to the formation of aggresome and clearance of unimported mitochondrial proteins. The autophagic gene *SESN2* is directly activated by the ISR network (*57*), whereas how the other autophagic genes are transcriptionally activated is yet to be determined. We found that mPOS also activates genes involved in proteasomal function. This is reminiscent of the UPRam mechanism that was recently reported in yeast when challenged by mitochondrial protein import stress (*10, 23*). Taken together, our data revealed that the skeletal muscle responds to mPOS by both suppressing protein synthesis, and by promoting protein sequestration and breakdown. These adaptive processes may collectively rebalance cytosolic proteostasis and reduce acute myofiber death. However, as a trade-off, chronic adaptations to mPOS lead to the overall reduction of protein content, which is ultimately manifested by myofiber shrinkage and muscle wasting (Fig. 7B).

Our results support an imbalance between mitochondrial protein load and import as a novel mechanism of muscle wasting. Although *ANT1* is a highly expressed gene, it is intriguing to observe that a two-fold increase in the expression of a single gene is sufficient to cause mPOS and drastic proteostatic adaptations in the cytosol. This highlights the vulnerability of the balance between mitochondrial protein loading and import capacity *in vivo*. Because unbalanced mitochondrial protein loading and import is expected to occur under many pathophysiological conditions, our finding could have potential implications for other myopathic diseases involving mitochondrial stress. For example, overexpression of nuclear genes encoding mitochondrial proteins may lead to preprotein overloading and saturation of the import machinery. Mutations in genes encoding the core protein import machinery may directly reduce the overall capacity of protein import relative to protein loading. IMM proteostatic stress and conditions affecting the generation and maintenance of membrane potential may indirectly reduce protein import efficiency. It is currently unknown whether it is the preprotein chaperoning and delivery systems in the cytosol or the level of import receptors/channels that limits the mitochondrial translocation of the excessive preproteins. As cytosolic aggresomes are readily detected, our data would also support that the cytosol has a limited capacity in degrading unimported mitochondrial preproteins in the skeletal muscle. In addition to the skeletal muscle, *ANT1* is also expressed in other tissues including the central nervous system. Despite the severe muscle wasting phenotype, we did not detect obvious phenotypes that may suggest neurodegeneration in the *ANT1*^*Tg*^/+ mice. The overall lifespan of the transgenic animals is unaffected. It is possible that the neuronal tissues have a better capacity than the skeletal muscle in handling mPOS. These tissues may possess a higher protein import capacity or a better controlled proteostatic network in the cytosol to efficiently degrade unimported mitochondrial preproteins.

We found that Ant1-induced mPOS leads to eIF2α phosphorylation and ISR activation. ISR activation has been previously documented in various models of mitochondrial stress (*33 37, 43–48*). What triggers eIF2α phosphorylation in *ANT1*^*Tg*^/+ muscles remains unsolved. Our RNA-Seq analysis did not detect the upregulation of anti-oxidant enzymes, which rules out the possibility that oxidative stress plays a major role in ISR activation. We found that mitochondrial respiration is moderately reduced in the *ANT1*^*Tg*^/+ muscles. Decreased mitochondrial respiration may result from an overall reduction in the import of OXPHOS components when cells are challenged by Ant1 overloading and the saturation of the protein import machinery. Given that more severe bioenergetic defect has very limited effect on muscle mass homeostasis during aging (*6–9*), it is unlikely that reduced ATP synthesis accounts for the severe muscle wasting phenotype in the *ANT1*^*Tg*^/+ mice. ISR activation independent of mitochondrial respiratory deficiency has been previously reported in a mouse model disrupted of *DARS2*, encoding a mitochondrial aspartyl-tRNA synthetase (*44*). Four stress-sensing kinases are known to promote eIF2α phosphorylation. These are HRI (EIF2AK1), PKR (EIF2AK2), PERK (EIF2AK3) and GCN2 (EIF2AK4) that are activated by heme deficiency/heavy metals, viral infection, ER stress and amino acid deficiency, respectively (*42*). Auwerx and coworkers showed that ISR activation by mitochondrial stressors in HeLa cells seems to be independent of the four eIF2α kinases (*48*). More recent studies showed that mitochondrial stress causes the release of the DELE1 protein, which in turn binds and activates the HRI kinase (*58, 59*). It would be of great interest in the future studies to determine whether Ant1 overloading induces eIF2α phosphorylation by the same mechanism, or by an independent pathway downstream of mPOS in the cytosol.

In addition to the activation of multiple pro-atrophic proteostatic pathways in *ANT1*^*Tg*^/+ muscles, it is relevant to note that Ant1-induced mPOS also activates *GADD45A* and *CDKN1A*, two additional downstream targets of the ISR network (*60*) (Supplemental Fig. 5J). *GADD45A* and *CDKN1A* encode the growth arrest and DNA-damage-inducible protein α, and the cyclin-dependent kinase inhibitor 1A (or p21) respectively. Both proteins promote muscle atrophy (*61–63*). *GADD45A* has also been shown to repress protein synthesis and activate proteolysis via an unknown mechanism. Thus, it is possible that *GADD45A* activation also contributes to the muscle wasting phenotype in the *ANT1*^*Tg*^/+ mice.

Our observation that a two-fold increase in *ANT1* expression is sufficient to cause severe muscle atrophy could have direct implications for human diseases such as FSHD and dilated cardiomyopathy that are associated with *ANT1* activation (*29, 30, 64, 65*). FSHD is primarily linked to the ectopic expression of *DUX4* in the skeletal muscle. It encodes a transcription factor that is not normally expressed in somatic tissue (*66*). In light of our study, it might be worth reexamining whether or not *ANT1* overexpression contributes to or modifies the course of this disease. Finally, our finding that mitochondrial protein import stress induces mPOS *in vivo* could also have implications for other clinical and subclinical conditions. Protein import is an elaborate process (*67*). Protein import efficiency is expected to be directly or indirectly affected in many human diseases (*68–72*). More broadly, aging is also associated with reduced protein import (*73*). Our finding may therefore help the understanding of how mitochondrial damage affects tissue homeostasis and function during aging and in import-related diseases.

## Materials and Methods

### Antibodies for western blot and immunofluorescence analysis

Antibodies used in this study included the Total OXPHOS Human WB Antibody Cocktail (#ab1104a1, Abcam), anti-Ant1 (#SAB2108761, Sigma), anti-Bax (#ab32503, Abcam), anti-eIF2 (#9722, Cell Signaling), anti-phospho-eIF2α (Ser51) (#3597, Cell Signaling), anti-4EBP1 (#9644, Cell Signaling), anti-Lamp2 (#GL2A7, Developmental Studies Hybridoma Bank, University of Iowa), anti-phospho-4EBP1 (#2855, Cell Signaling), anti-p62 (#23214, Cell Signaling), anti-rpS6 (#ab40820, Abcam), anti-phospho-rpS6 (#ab65748, Abcam), anti-ubiquitin (#701339, Thermo Scientific). Total proteins were stained with REVERT Total Protein Stain (#926-11011, LI-COR). For western blot, band intensities were quantified by the LI-COR Odyssey Fc Imaging system and normalized against total protein staining.

### Generation of *ANT1*-transgenic mice

The Ant1 transgene was prepared by recombineering according to Lee et al.(*74*). Briefly, genomic sequences corresponding to Ant1 5’ upstream and 3’ downstream genomic sequence were prepared by PCR using primer pairs ret5F (5’-GTCGAATTC GTATATAAATAAATAAAAGAAAG) and ret5R (5’-GTCAGACGTC CAATGTTGCTACTTAAACACTCTTG) and ret3F (5’ GTCA GACGTC CCTTGAGAACTAACACAGAGCAG) and ret3R (5’ GTCAAAGCTT GGTGATATGGGGACAGGAAGGAG). The 5’ and 3’ mini-arms were digested with *Eco*RI and *Aat*II, and *Hin*dIII and *Aat*II, respectively, and then inserted into pSK+. The mini-arm vector was then digested with *Aat*II, treated with phosphatase, and then electroporated into EL350 together with a BAC clone, RP24-108A1, which contains the entire Ant1 genomic sequence, to retrieve Ant1 genomic sequence from the BAC by gap repair. The retrieved Ant1 genomic sequence is 13.7 kb in size containing all 4 exons together with 4.2 kb and 4.96 kb of 5’ up- and 3’ down-stream sequences, respectively. The Ant1 transgene fragment was released from the plasmid by *Not*I and *Kpn*I digestion, fractionated by agarose gel electrophoresis, column purified and resuspended in 10mM Tris, pH8.0.

*ANT1*-transgenic mice were generated according to Nagy et al.(*75*). Briefly, four to six week old C57BL6j females were superovulated by first administration of 5 IU of pregnant mare serum (PMS) and 44 to 46 hour later with 5 IU human chorionic gonadotropins (HCG). The females are then mated with C57BL6j stud males. Fertilized one-cell embryos were collected from the oviduct the next day and maintained in KSOM medium in an air/CO2 incubator at 37°C until use. Purified Ant1 genomic fragment were diluted to approximately 1 ng/μl and then microinjected into the pronuclei of the one-cell embryos. Injected embryos were implanted into the oviducts of 0.5 days *post coitium* pseudopregnant females for subsequent development *in utero*. Transgenic mice were identified by PCR using genomic DNA prepared from ear notch as template. Two primer pairs specific to the 5’ and 3’ end of the Ant1 transgene were used for genotyping. Primer pair BHtoRI (5’-GGATCCCCCGGGCTGCAGGAATTC) and Ant R5 (5’-CAATGTTGCTACTTAAACACTCTTG), corresponds to the 5’ will identify a fragment of 515 bp, and a second pair of primers, XhotoHind (5’-GAGGTCGACGGTATCGATAAGCTTG) and AntR3F (5’-CTTTCCTGGACCCCTGTAAGCTTG) will detect a fragment of 432 bp specific to the 3’ end of the transgene. Founder animals were then mated with C57BL6NTac mice to establish and maintain the transgenic line. All the animal experiments have been approved by the Institutional Animal Care and Use Committee (IACUC) of the State University of New York Upstate Medical University. The experiments were performed using age-matched and mostly littermate controls. The animals were fed with a regular chow diet *ad libitum* and were housed at an ambient temperature. Food and water were placed at a low and reachable position for the *ANT1*-transgenic mice over one-year-old.

### Lean mass, home cage activity and exercise endurance tests

Lean mass and home cage activity were measured by quantitative magnetic resonance and Promethion metabolic caging respectively at Vanderbilt University Mouse Metabolic Phenotyping Center. Exercise endurance was measured with an Exer 3/6 Treadmill (Columbus Instruments) by determining the maximum distance and speed of the animals before exhaustion. The animals were trained for at least two sessions before the tests.

### Mitochondrial isolation and bioenergetic assay

Unanesthetized mice were decapitated using a guillotine. Muscle mitochondria were isolated according to Garcia-Cazabin et al. (*76*). Oxygen consumption rates were measured using an Oxygraph Plus oxygen electrode (Hansatech Instruments Ltd), with 150 μg mitochondria, 5 mM glutamate, 2.5 mM malate, 150 μM ADP. Oligomycin (5 μg/ml) was added to established state-4 respiration. Respiratory control ratio was established by dividing state-3 respiratory rate with oligomycin-inhibited respiratory rate.

### Muscle histology and immunohistochemistry

Standard procedures were followed for H&E, Trichrome, NADH, SDH and COX activity staining, and for the determination of lesser diameter and variability coefficient of muscle fibers (*77*). For immunofluorescence microscopy, muscle sections of 10 μm thickness were fixed and permeabilized with 100% methanol at −20°C for 15 minutes, rinsed in PBS, treated in blocking buffer (1XPBS/5% Normal Goat Serum/0.3% Triton X-100) for one hour at room temperature, followed by incubation with anti-Lamp2 antibody at 4 °C overnight. The specimens were then washed in 1XPBS/0.1% Tween 20 before being probed with an Alexa Fluor 488-conjugated anti-rat IgG (H+L) antibody for one hour at room temperature. After washing with 1XPBS/0.1% Tween 20, the tissue samples were mounted with ProLong Diamond Antifade Mountant with DAPI (#P36962, Invitrogen), and were visualized using a Leica SP8 confocal microscope.

### Electron microscopy

Fresh muscle samples were fixed in 4% glutaraldehyde/0.1 M cacodylate buffer, pH7.2 at room temperature for two hours, and postfixed with 1% osmium tetroxide/0.1 M cacodylate buffer, pH7.2 at room temperature for one hour. The specimens were dehydrated in 50%, 70%, 90%, 100% ethanol and propylene oxide, before being embedded in Luft’s Araldite 502 embedding medium (Electron Microscopy Sciences, Hatfield, PA) and cut into thin sections. The ultrathin sections were then stained with ethanolic uranyl acetate and Reynold’s lead citrate (Polysciences). The samples were examined with a JOEL JEM1400 transmission electron microscope and images were acquired with a Gaten DAT-832 Orius camera.

### RNA-Seq analysis

Total RNA was extracted from snap-frozen quadriceps muscles (n=4/genotype/sex) using RNeasy mini kit (Qiagen). The quality of total RNA was validated by Bioanalyzer (Agilent Technologies). Approximately 1 µg of RNA per sample was used to construct the cDNA library using TruSeq stranded mRNA library prep kit (Illumina). The cDNA libraries were quantitated using KAPA library quantification kit for Illumina platforms (Kapa Biosystems). The individual indexed libraries were diluted to 4 nM and pooled in equal quantity, denatured before loading onto the Illumina NextSeq 500. The sequencing was run as paired-end reads (2 × 75 bp per read) with a targeted depth of 60 million paired-end reads per sample. Sequence samples were then aligned and quantified with the Salmon algorithm (*78*) using default parameters in RaNA-seq. Differential expression analysis was performed using a one-way analysis of variance (ANOVA) to test the main effect of genotype using both limma (*79*) following quantile normalization and DESeq2 (*80*). This yielded a total of 2076 and 2654 significant genes at a *q*-value cutoff of 0.01, respectively. These lists were subjected to volcano plot visualization and hierarchical clustering analysis. In addition to single findings, we also tested for Gene Set Enrichment Analysis (GSEA) using a rapid modified algorithm (*81*), and displayed the top 20 Pathways and Gene Ontologies in graphical and tabular form.

### Determination of protein/DNA ratio

DNA and proteins were extracted from approximately 30 of quadriceps muscles using AllPrep DNA/RNA/Protein Mini kit (#80004, Qiagen). DNA concentrations were determined by Thermo Scientific NanoDrop 2000c spectrophotometer. Protein concentrations were determined by Bradford protein assay (#**5000006, Bio-Rad)**. Relative protein contents in the muscles were calculated after normalizing by total amount of DNA.

### Proteasomal activity assay

Quadriceps muscle samples (1 mg) were sonicated in PBS plus EDTA (5 mM, pH 7.4) buffer three times for 5 seconds with 25 seconds intervals on ice. After the removal of cell debris by centrifugation, protein concentration of the soluble fractions was determined by Bradford assay. 10 μg of the muscle lysates were used for determining proteasome-associated chymotrypsin-like activity with the Proteasome-Glo^TM^ Assay Systems (#G8531, Promega). Luminescence signals were detected with the SpectraMax i3x Multi-Mode Microplate Reader (Molecular Devices) between 10–30 minutes at room temperature after the addition of luminogenic substrates. The chymotrypsin-like activity were calculated by subtracting MG132 (50 μM)-inhibited activities from total activity.

## Acknowledgments

We thank David Mitchell for help on electron microscopy, Siu-Pok Yee (University of Connecticut) for the generation of the *ANT1*-transgenic mice, Don Henderson (University of Rochester) and Christopher Turner for muscle histology, Jushuo Wang for confocal microscopy, SUNY Upstate Medical University Department of Laboratory Animal Resources for animal care, Vanderbilt University Mouse Metabolic Phenotyping Center (MMPC) for mouse phenotyping, Michael Sherman (Boston University) for providing plasmids, and the Chen laboratory members for comments on the manuscript. This work was supported by the NIH grants AG061204 and AG047400 to X.J.C.. X.W., X.J.C. and R.T. performed the experiments. F.M. processed and analyzed the RNA-Seq data. X.W. and X.J.C. wrote the manuscript. The authors declare no competing financial interests. All raw FastQ files and processed read alignment count data from the RNA-Seq experiments have been deposited into the NCBI Gene Expression Omnibus/Sequence Read Archive (accession number: GSE135584).

## Supplementary Materials

### This PDF file includes

Supplemental Figure 1 to 6.

Captions for Movies 1 to 5.

Captions for Supplemental Table 1 to 3.

### Other Supplementary Materials for this manuscript include the following

Supplemental Movies 1 to 5 Supplemental Tables 1 to 3

**Supplementary Figure 1.**
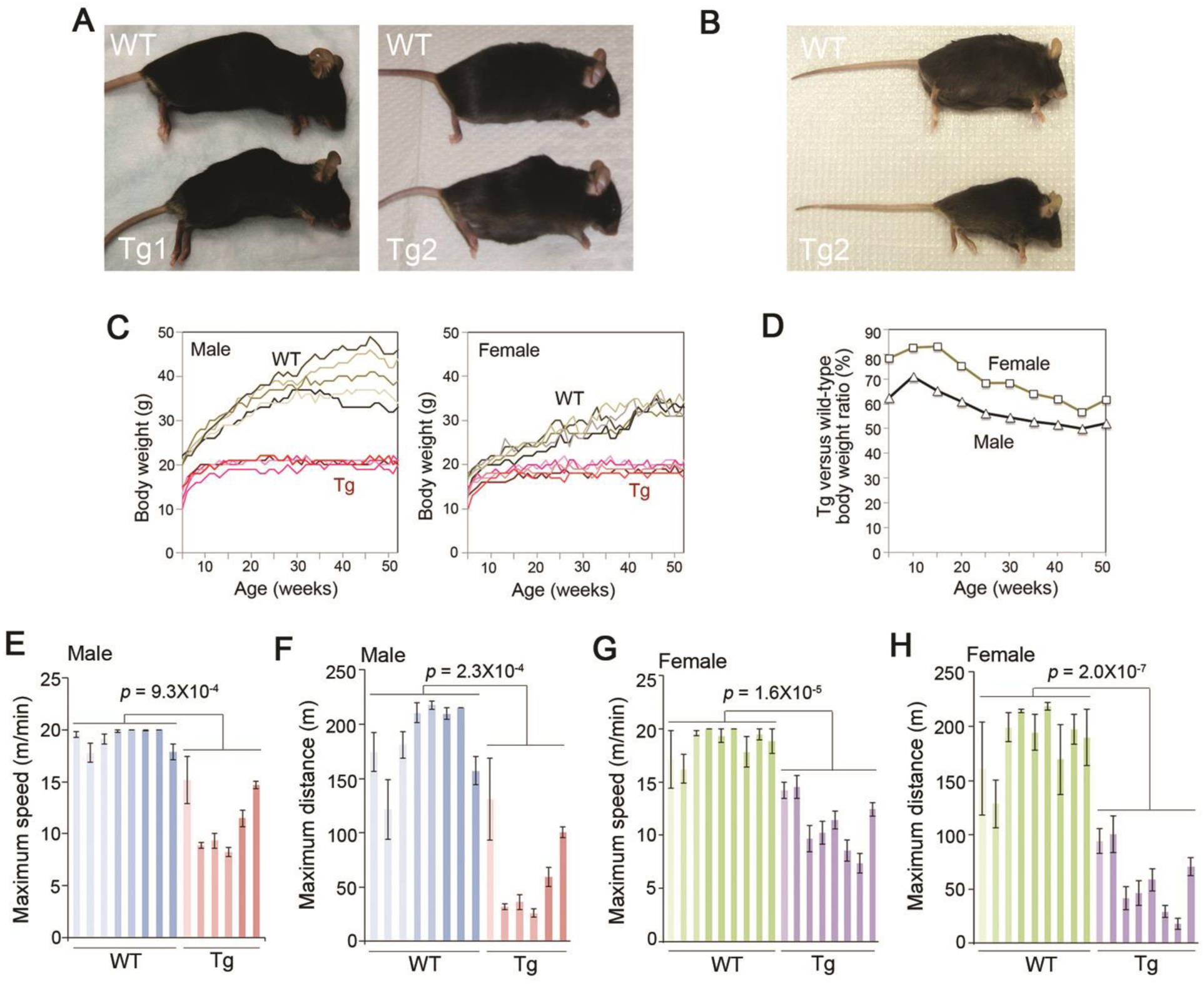
Body morphology, weight and exercise endurance of *ANT1*^*Tg*^/+ mice compared with wild-type controls. (**A**) Kyphotic phenotype in two independent *ANT1*^*Tg*^/+ lines at the age of 3 (left) and 6 (right) months respectively. (**B**) Kyphotic phenotype of *ANT1*^*Tg*^/+ mice at 17 months. (**C**) and (**D**) Body weight of *ANT1*^*Tg*^/+ mice (n=5/genotype/sex). (**E-H**) Exercise endurance of *ANT1*^*Tg*^/+ mice compared with littermate controls (6.5-8.5 months old; n=6-9/genotype/sex). The maximum speed and distance of wild-type mice were not tested beyond 20 m/min and 200 m respectively. Compared with control mice, the *ANT1*^*Tg*^/+ mice have significant decreases in all the activity parameters measured, except for female *ANT1*^*Tg*^/+ mice that have only a marginally reduced walk speed. m, meter. Error bars represent S.E.M. of 4-6 tests for each mouse. *P* values were calculated by unpaired Student’s *t* test.

**Supplementary Figure 2.**
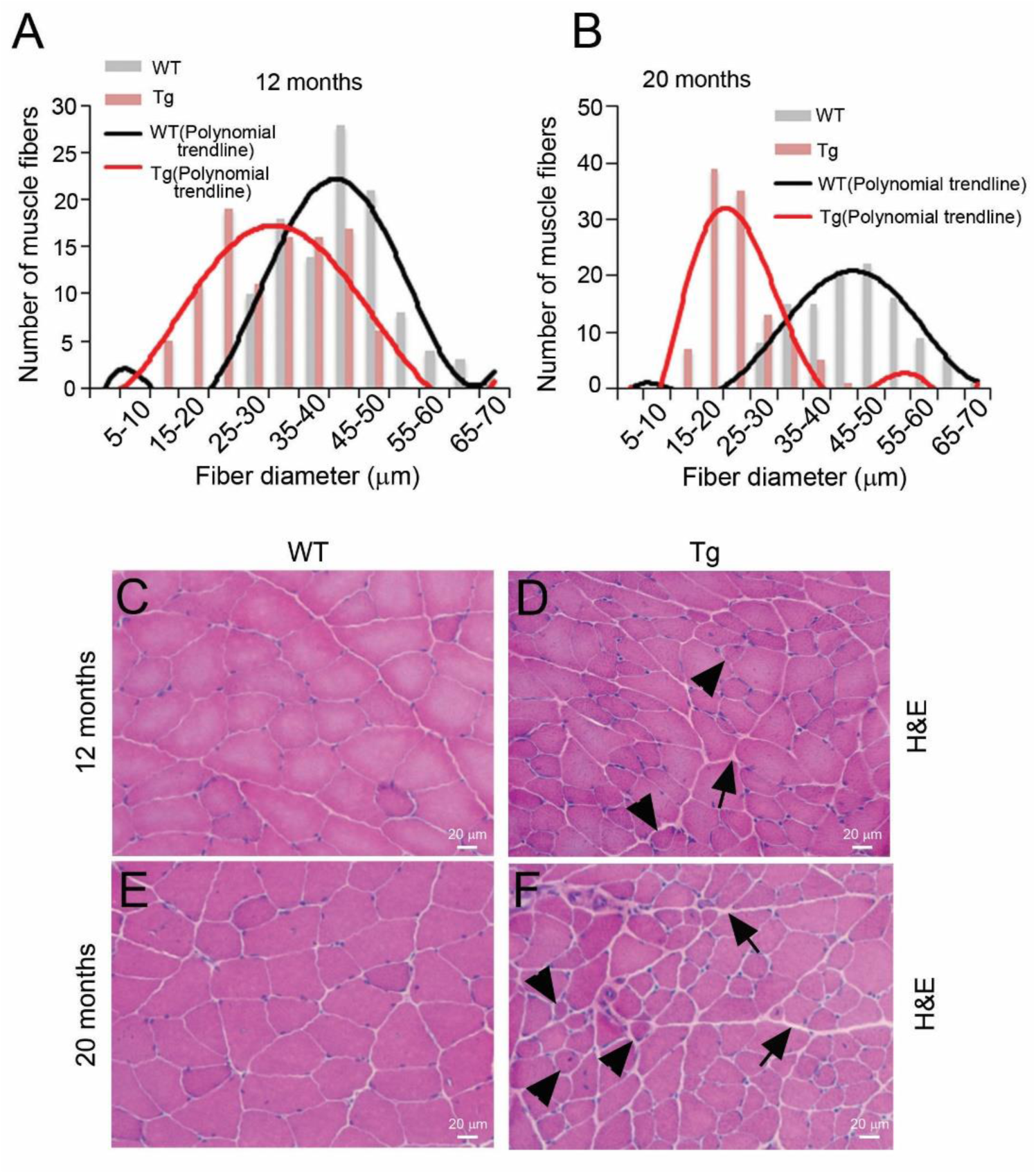
Histological analysis of *ANT1*^*Tg*^/+ muscles. (**A**) and (**B**) Muscle fiber size distribution at the age of 12 and 20 months. (**C**) – (**F**), hematoxylin and eosin (H&E) staining of *ANT1*^*Tg*^/+ quadriceps muscles compared with wild type controls. Arrows and arrow heads denote endomyosium and round- and angular-shaped small myofibers, respectively.

**Supplementary Figure 3.**
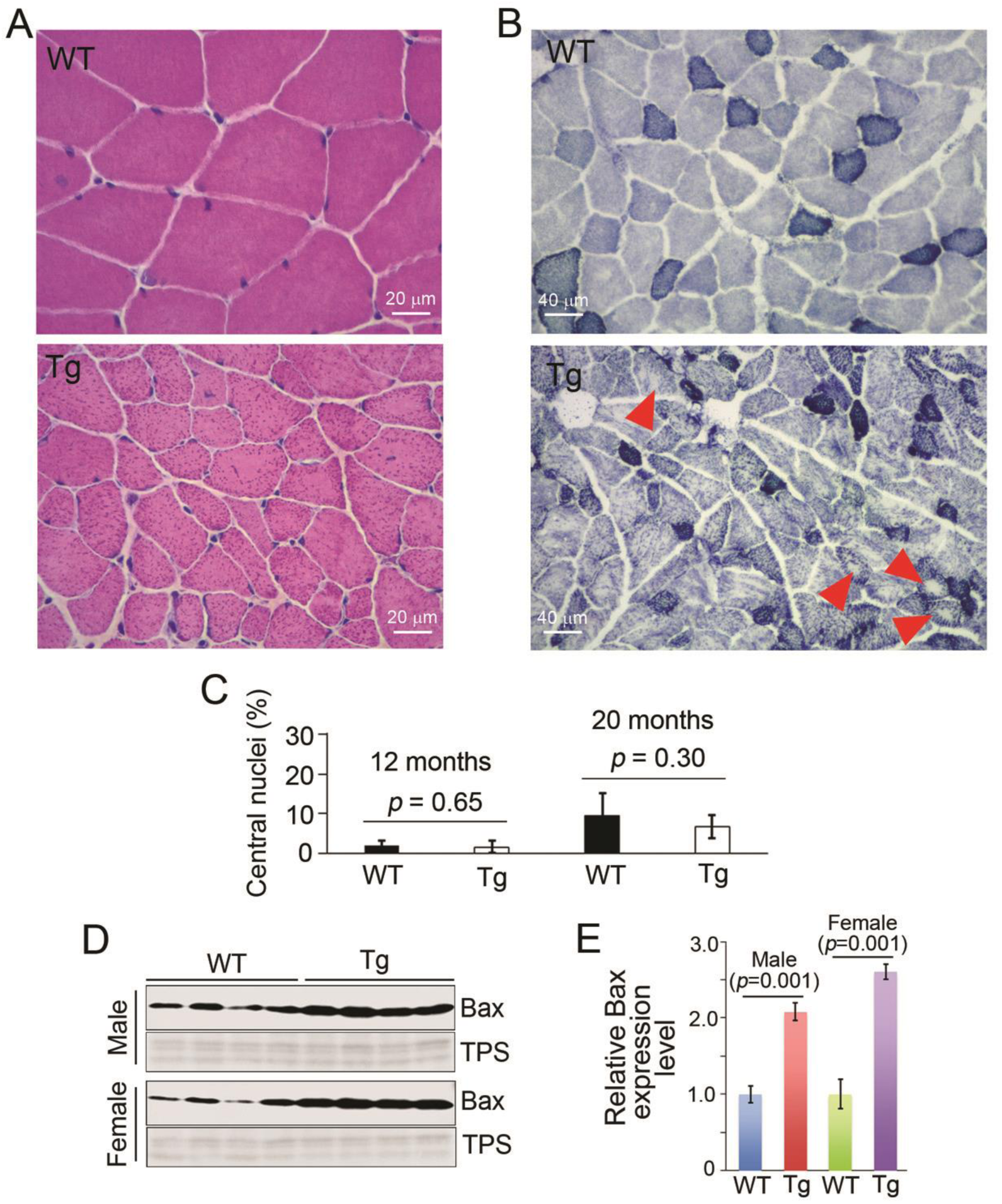
Basophilic structures, “moth-eaten” myofibers, nuclear positioning and *BAX* expression in *ANT1*^*Tg*^/+ muscles. (**A**) H&E staining of quadriceps showing basophilic stippling in a 20-month-old female *ANT1*^*Tg*^/+ mouse. (**B**) NADH staining of quadriceps showing “moth-eaten” in a 20-month-old female *ANT1*^*Tg*^/+ mouse. (**C**) Ratios of myofibers with central nuclei in *ANT1*^*Tg*^/+ and control mice at the age of 12 and 20 months. Totally 296 - 451 myofibers from 5-8 different fields were counted at each time point. (**D**) and (**E**) Increased steady state levels of Bax (n=4/genotype/sex), as measured by western blot. TPS, total protein staining.

**Supplementary Figure 4.**
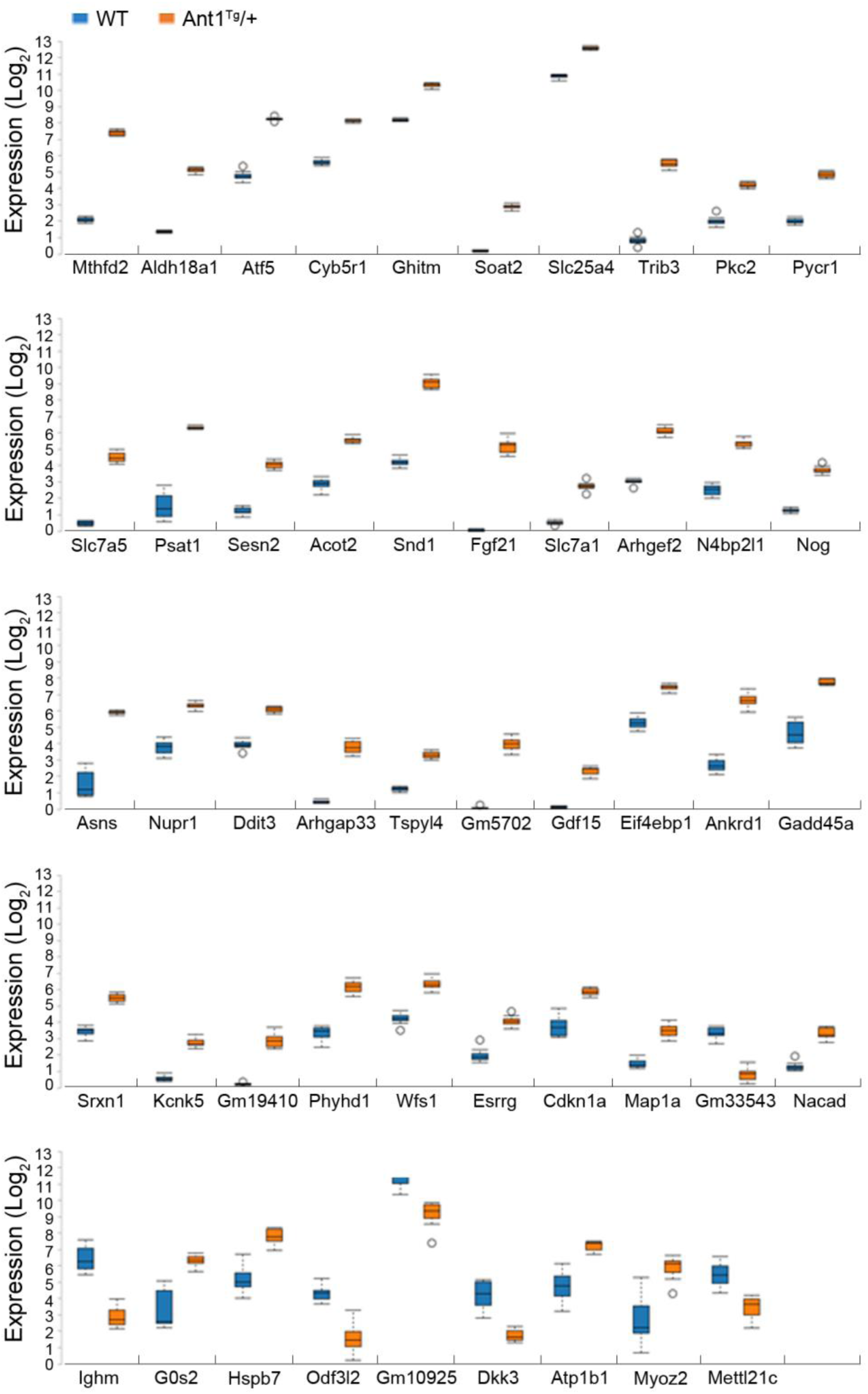
Whisker-box plots of 48 genes with 4-fold changes in *ANT1*^*Tg*^/+ mice versus WT mice, at FDR < 0.01, with *ANT1* (Slc25a4) included for comparative purposes.

**Supplementary Figure 5.**
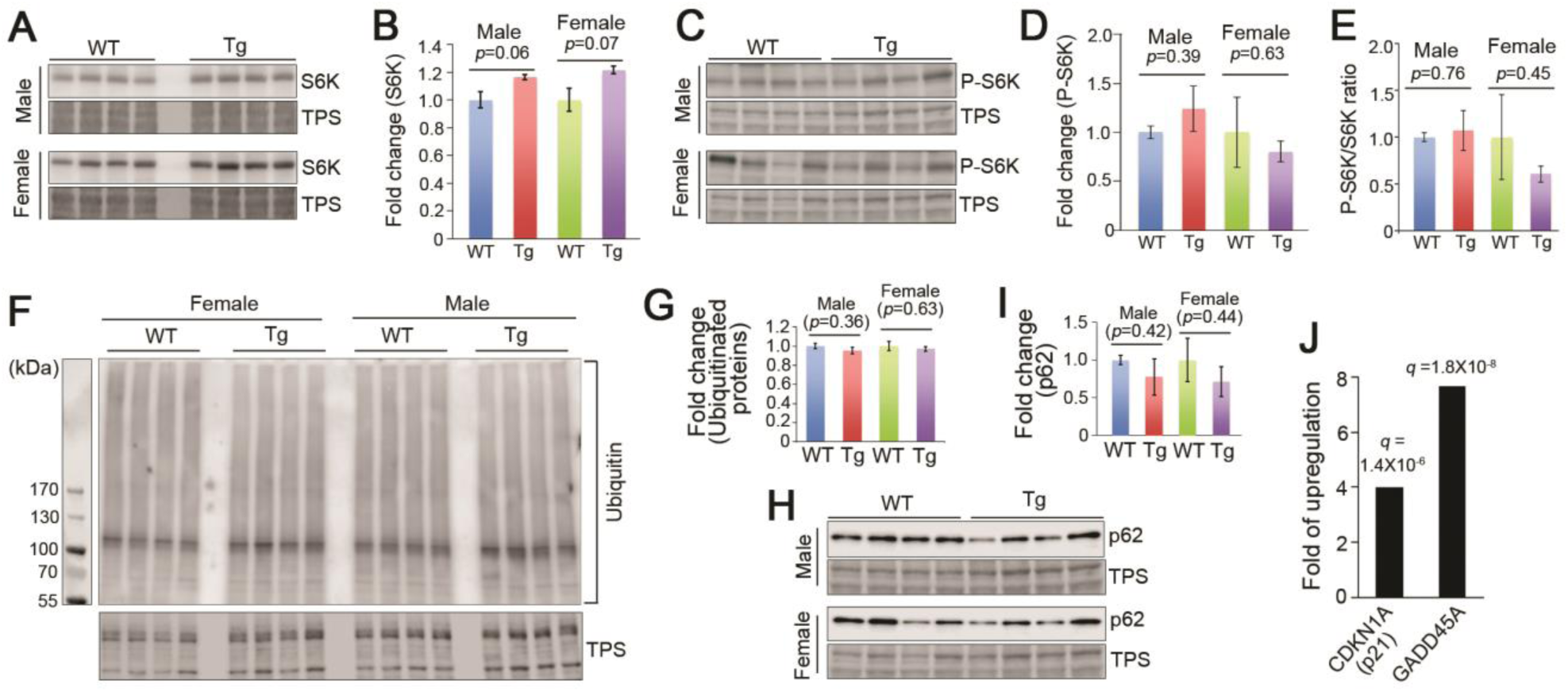
RpsS6 kinase phosphorylation, protein ubiquitination, p62 expression, and *GADD45A* and *CDKN1A* activation in *ANT1*^*Tg*^/+ mice at 6 months old (n=4/genotype/sex) (**A &B**) Western-blot showing the levels of RpsS6 kinase (S6K). (**C&D**) Western-blot showing the levels of phosphorylated S6K (P-S6K). (**E**) P-S6K/S6K ratio. (**F**) Western blot showing ubiquitinated proteins using an anti-ubiquitin antibody. The relative ubiquitination levels were quantified and shown in (**G**). (**H**&**I**) Steady state levels of p62, as determined by western blot. (**J**) Transcriptional upregulation of *GADD45A* and *CDKN1A* in *ANT1^Tg^/+* muscles.

**Supplementary Figure 6.**
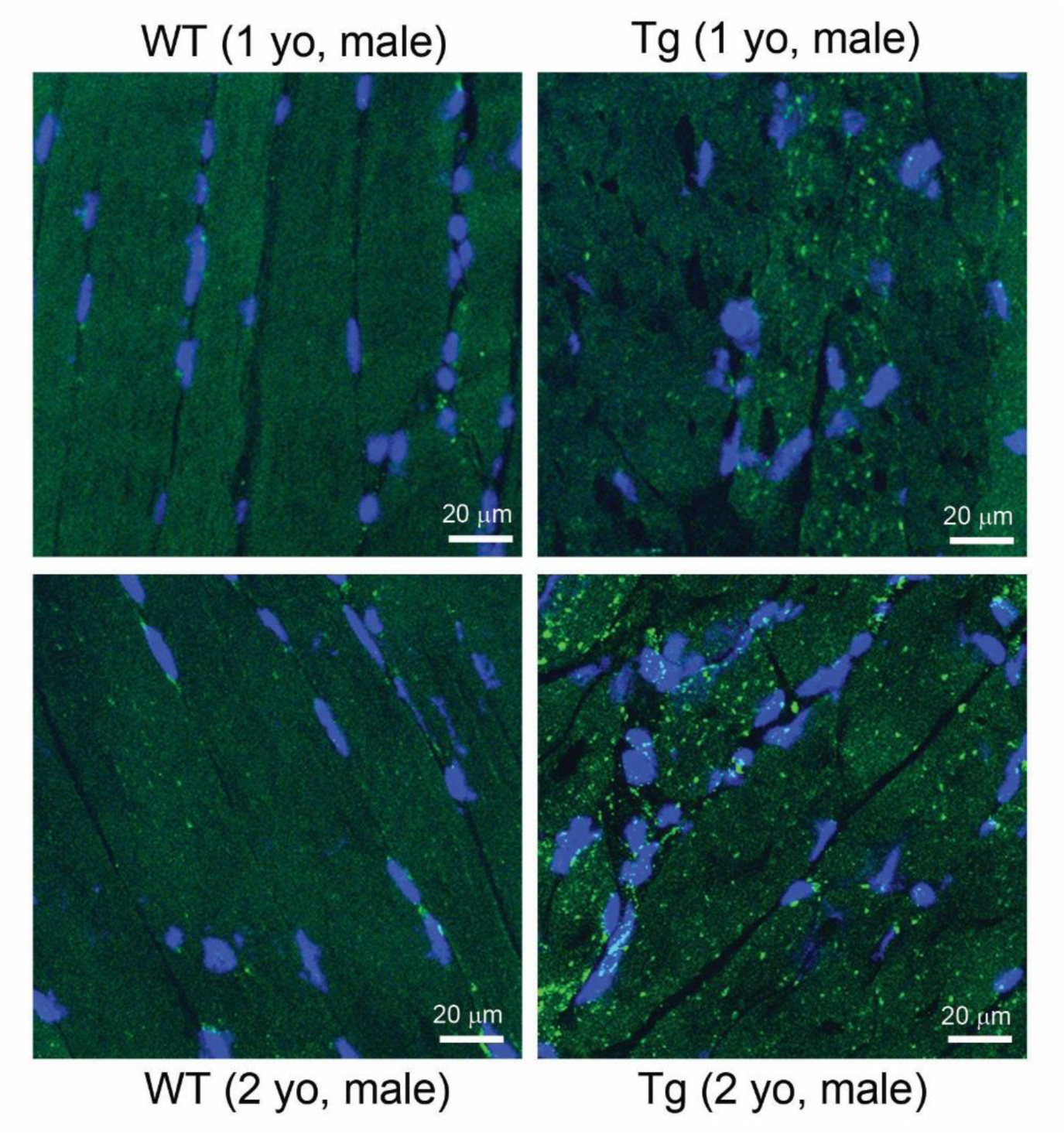
Confocal microscopy showing Lamp2-positive lysosomal and lysosome-associated cellular structures in quadriceps muscles from *ANT1*^*Tg*^/+ and control mice. Longitudinal sections were examined in one- and two-year-old male mice. Blue, DAPI-stained nuclei; green, Lamp2-positive structures.

### Supplemental movies

**Movie S1** – Gait abnormalities of an *ANT1*^*Tg*^/+ male mouse at the age of 7 months, with a wild-type littermate as control.

**Movie S2 –** Gait abnormalities of an *ANT1*^*Tg*^/+ male mouse at the age of 12 months, with a wild-type littermate as control. The same pair of mice were filmed as in Supplemental movie 1.

**Movie S3 –** Gait abnormalities of an *ANT1*^*Tg*^/+ male mouse at the age of 19 months, with a wild-type littermate as control. The same pair of mice were filmed as in Supplemental movie 1.

**Movie S4 –** 3D movie showing a quadriceps section (10 m) from a 12-month old wild-type female mouse, stained with Lamp2 for lysosome and lysosome-derived structures (green), and DAPI for the nuclei (blue). The data are shown in a separate MP4 file.

**Movie S5 –** 3D movie showing a quadriceps section (10 m) from a 12-month old female *ANT1*^*Tg*^/+ mouse, stained with Lamp2 for lysosome and lysosome-derived structures (green), and DAPI for the nuclei (blue). The data are shown in a separate MP4 file.

### Supplemental Table 1 to 3

**Supplemental Table 1**. List of genes detected by RNA-Seq of quadriceps muscles from *ANT1*^*Tg*^/+ and wild-type mice. The data are shown in a separate Excel file.

**Supplemental Table 2**. List of genes whose transcription is changed by over 4 fold in *ANT1*^*Tg*^/+ muscles. The data are shown in a separate Excel file.

**Supplemental Table 3**. GSEA pathways. The data are shown in a separate Excel file.

## Notes

### Competing Interest Statement

The authors have declared no competing interest.

## References

1. J. A. Batsis, T. A. Mackenzie, L. K. Barre, F. Lopez-Jimenez, S. J. Bartels, Sarcopenia, sarcopenic obesity and mortality in older adults: results from the National Health and Nutrition Examination Survey III. Eur J Clin Nutr 68, 1001–1007 (2014).

2. S. Cohen, J. A. Nathan, A. L. Goldberg, Muscle wasting in disease: molecular mechanisms and promising therapies. Nat Rev Drug Discov 14, 58–74 (2015).

3. H. Chen et al., Mitochondrial fusion is required for mtDNA stability in skeletal muscle and tolerance of mtDNA mutations. Cell 141, 280–289 (2010).

4. R. Calvani et al., Mitochondrial pathways in sarcopenia of aging and disuse muscle atrophy. Biol Chem 394, 393–414 (2013).

5. V. Romanello, M. Sandri, Mitochondrial Quality Control and Muscle Mass Maintenance. Front Physiol 6, 422 (2016).

6. M. S. Lustgarten et al., MnSOD deficiency results in elevated oxidative stress and decreased mitochondrial function but does not lead to muscle atrophy during aging. Aging Cell 10, 493–505 (2011).

7. M. S. Lustgarten et al., Conditional knockout of Mn-SOD targeted to type IIB skeletal muscle fibers increases oxidative stress and is sufficient to alter aerobic exercise capacity. Am J Physiol Cell Physiol 297, C1520–1532 (2009).

8. Y. Zhang et al., CuZnSOD gene deletion targeted to skeletal muscle leads to loss of contractile force but does not cause muscle atrophy in adult mice. FASEB J 27, 3536–3548 (2013).

9. R. M. Morrow et al., Mitochondrial energy deficiency leads to hyperproliferation of skeletal muscle mitochondria and enhanced insulin sensitivity. Proc Natl Acad Sci U S A 114, 2705–2710 (2017).

10. U. Topf, L. Wrobel, A. Chacinska, Chatty Mitochondria: Keeping Balance in Cellular Protein Homeostasis. Trends Cell Biol, (2016).

11. L. P. Coyne, X. J. Chen, mPOS is a novel mitochondrial trigger of cell death - implications for neurodegeneration. FEBS Lett 592, 759–775 (2018).

12. E. I. Rugarli, T. Langer, Mitochondrial quality control: a matter of life and death for neurons. EMBO J 31, 1336–1349 (2012).

13. N. U. Naresh, C. M. Haynes, Signaling and Regulation of the Mitochondrial Unfolded Protein Response. Cold Spring Harb Perspect Biol 11, (2019).

14. R. J. Hunt et al., Mitochondrial stress causes neuronal dysfunction via an ATF4-dependent increase in L-2-hydroxyglutarate. J Cell Biol 218, 4007–4016 (2019).

15. J. R. Veatch, M. A. McMurray, Z. W. Nelson, D. E. Gottschling, Mitochondrial dysfunction leads to nuclear genome instability via an iron-sulfur cluster defect. Cell 137, 1247–1258 (2009).

16. X. Wang, The expanding role of mitochondria in apoptosis. Genes Dev 15, 2922–2933 (2001).

17. J. Nunnari, A. Suomalainen, Mitochondria: in sickness and in health. Cell 148, 1145–1159 (2012).

18. A. P. West et al., Mitochondrial DNA stress primes the antiviral innate immune response. Nature 520, 553–557 (2015).

19. Q. Zhang et al., The Mitochondrial Unfolded Protein Response Is Mediated Cell-Non-autonomously by Retromer-Dependent Wnt Signaling. Cell 174, 870–883 e817 (2018).

20. X. Wang, X. J. Chen, A cytosolic network suppressing mitochondria-mediated proteostatic stress and cell death. Nature 524, 481–484 (2015).

21. C. U. Martensson et al., Mitochondrial protein translocation-associated degradation. Nature 569, 679–683 (2019).

22. H. Weidberg, A. Amon, MitoCPR-A surveillance pathway that protects mitochondria in response to protein import stress. Science 360, (2018).

23. L. Wrobel et al., Mistargeted mitochondrial proteins activate a proteostatic response in the cytosol. Nature 524, 485–488 (2015).

24. F. Boos et al., Mitochondrial protein-induced stress triggers a global adaptive transcriptional programme. Nat Cell Biol 21, 442–451 (2019).

25. A. M. Nargund, M. W. Pellegrino, C. J. Fiorese, B. M. Baker, C. M. Haynes, Mitochondrial import efficiency of ATFS-1 regulates mitochondrial UPR activation. Science 337, 587–590 (2012).

26. Y. F. Lin, C. M. Haynes, Metabolism and the UPR(mt). Mol Cell 61, 677–682 (2016).

27. D. Poveda-Huertes et al., An Early mtUPR: Redistribution of the Nuclear Transcription Factor Rox1 to Mitochondria Protects against Intramitochondrial Proteotoxic Aggregates. Mol Cell 77, 180–188 e189 (2020).

28. Y. Liu et al., Mitochondrial carrier protein overloading and misfolding induce aggresomes and proteostatic adaptations in the cytosol. Mol Biol Cell 30, 1272–1284 (2019).

29. D. Gabellini, M. R. Green, R. Tupler, Inappropriate gene activation in FSHD: a repressor complex binds a chromosomal repeat deleted in dystrophic muscle. Cell 110, 339–348 (2002).

30. D. Laoudj-Chenivesse et al., Increased levels of adenine nucleotide translocator 1 protein and response to oxidative stress are early events in facioscapulohumeral muscular dystrophy muscle. J Mol Med 83, 216–224 (2005).

31. D. Gabellini et al., Facioscapulohumeral muscular dystrophy in mice overexpressing *FRG1*. Nature 439, 973–977 (2006).

32. M. D. Brand et al., The basal proton conductance of mitochondria depends on adenine nucleotide translocase content. Biochem J 392, 353–362 (2005).

33. F. Ishikawa et al., Gene expression profiling identifies a role for CHOP during inhibition of the mitochondrial respiratory chain. J Biochem 146, 123–132 (2009).

34. H. Tyynismaa et al., Mitochondrial myopathy induces a starvation-like response. Hum Mol Genet 19, 3948–3958 (2010).

35. J. Nikkanen et al., Mitochondrial DNA Replication Defects Disturb Cellular dNTP Pools and Remodel One-Carbon Metabolism. Cell Metab 23, 635–648 (2016).

36. X. R. Bao et al., Mitochondrial dysfunction remodels one-carbon metabolism in human cells. Elife 5, (2016).

37. I. Kuhl et al., Transcriptomic and proteomic landscape of mitochondrial dysfunction reveals secondary coenzyme Q deficiency in mammals. Elife 6, (2017).

38. M. Ost et al., Muscle mitohormesis promotes cellular survival via serine/glycine pathway flux. FASEB J 29, 1314–1328 (2015).

39. S. Carra et al., The growing world of small heat shock proteins: from structure to functions. Cell Stress Chaperones 22, 601–611 (2017).

40. A. Mogk, C. Ruger-Herreros, B. Bukau, Cellular Functions and Mechanisms of Action of Small Heat Shock Proteins. Annu Rev Microbiol 73, 89–110 (2019).

41. J. Kirstein-Miles, A. Scior, E. Deuerling, R. I. Morimoto, The nascent polypeptide-associated complex is a key regulator of proteostasis. EMBO J 32, 1451–1468 (2013).

42. K. Pakos-Zebrucka et al., The integrated stress response. EMBO Rep 17, 1374–1395 (2016).

43. J. M. Silva, A. Wong, V. Carelli, G. A. Cortopassi, Inhibition of mitochondrial function induces an integrated stress response in oligodendroglia. Neurobiol Dis 34, 357–365 (2009).

44. S. A. Dogan et al., Tissue-specific loss of DARS2 activates stress responses independently of respiratory chain deficiency in the heart. Cell Metab 19, 458–469 (2014).

45. N. A. Khan et al., mTORC1 Regulates Mitochondrial Integrated Stress Response and Mitochondrial Myopathy Progression. Cell Metab 26, 419–428 e415 (2017).

46. O. Zurita Rendon, E. A. Shoubridge, LONP1 Is Required for Maturation of a Subset of Mitochondrial Proteins, and Its Loss Elicits an Integrated Stress Response. Mol Cell Biol 38, (2018).

47. B. M. Baker, A. M. Nargund, T. Sun, C. M. Haynes, Protective coupling of mitochondrial function and protein synthesis via the eIF2alpha kinase GCN-2. PLoS Genet 8, e1002760 (2012).

48. P. M. Quiros et al., Multi-omics analysis identifies ATF4 as a key regulator of the mitochondrial stress response in mammals. J Cell Biol 216, 2027–2045 (2017).

49. P. Bonaldo, M. Sandri, Cellular and molecular mechanisms of muscle atrophy. Dis Model Mech 6, 25–39 (2013).

50. M. J. Kim et al., SESN2/sestrin2 suppresses sepsis by inducing mitophagy and inhibiting NLRP3 activation in macrophages. Autophagy 12, 1272–1291 (2016).

51. A. Kumar, C. Shaha, SESN2 facilitates mitophagy by helping Parkin translocation through ULK1 mediated Beclin1 phosphorylation. Sci Rep 8, 615 (2018).

52. S. Jiang et al., Starch binding domain-containing protein 1/genethonin 1 is a novel participant in glycogen metabolism. J Biol Chem 285, 34960–34971 (2010).

53. T. Nakazawa et al., Emerging roles of ARHGAP33 in intracellular trafficking of TrkB and pathophysiology of neuropsychiatric disorders. Nat Commun 7, 10594 (2016).

54. M. Krendel, F. T. Zenke, G. M. Bokoch, Nucleotide exchange factor GEF-H1 mediates cross-talk between microtubules and the actin cytoskeleton. Nat Cell Biol 4, 294–301 (2002).

55. S. A. Eisler et al., A Rho signaling network links microtubules to PKD controlled carrier transport to focal adhesions. Elife 7, (2018).

56. C. A. Juliana et al., ATF5 regulates beta-cell survival during stress. Proc Natl Acad Sci U S A 114, 1341–1346 (2017).

57. A. A. Garaeva, I. E. Kovaleva, P. M. Chumakov, A. G. Evstafieva, Mitochondrial dysfunction induces SESN2 gene expression through Activating Transcription Factor 4. Cell Cycle 15, 64–71 (2016).

58. X. Guo et al., Mitochondrial stress is relayed to the cytosol by an OMA1-DELE1-HRI pathway. Nature 579, 427–432 (2020).

59. E. Fessler et al., A pathway coordinated by DELE1 relays mitochondrial stress to the cytosol. Nature 579, 433–437 (2020).

60. C. M. Adams, S. M. Ebert, M. C. Dyle, Role of ATF4 in skeletal muscle atrophy. Curr Opin Clin Nutr Metab Care 20, 164–168 (2017).

61. S. M. Ebert et al., Stress-induced skeletal muscle Gadd45a expression reprograms myonuclei and causes muscle atrophy. J Biol Chem 287, 27290–27301 (2012).

62. K. S. Bongers et al., Skeletal muscle denervation causes skeletal muscle atrophy through a pathway that involves both Gadd45a and HDAC4. Am J Physiol Endocrinol Metab 305, E907–915 (2013).

63. D. K. Fox et al., p53 and ATF4 mediate distinct and additive pathways to skeletal muscle atrophy during limb immobilization. Am J Physiol Endocrinol Metab 307, E245-261 (2014).

64. E. Kim et al., ZNF555 protein binds to transcriptional activator site of 4qA allele and ANT1: potential implication in Facioscapulohumeral dystrophy. Nucleic Acids Res 43, 8227–8242 (2015).

65. A. Dorner, K. Schulze, U. Rauch, H. P. Schultheiss, Adenine nucleotide translocator in dilated cardiomyopathy: pathophysiological alterations in expression and function. Mol Cell Biochem 174, 261–269 (1997).

66. J. M. Statland, R. Tawil, Facioscapulohumeral muscular dystrophy: molecular pathological advances and future directions. Curr Opin Neurol 24, 423–428.

67. A. Chacinska, C. M. Koehler, D. Milenkovic, T. Lithgow, N. Pfanner, Importing mitochondrial proteins: machineries and mechanisms. Cell 138, 628–644 (2009).

68. C. M. Koehler et al., Human deafness dystonia syndrome is a mitochondrial disease. Proc Natl Acad Sci U S A 96, 2141–2146 (1999).

69. Y. Kang et al., Sengers Syndrome-Associated Mitochondrial Acylglycerol Kinase Is a Subunit of the Human TIM22 Protein Import Complex. Mol Cell 67, 457–470 e455 (2017).

70. M. Vukotic et al., Acylglycerol Kinase Mutated in Sengers Syndrome Is a Subunit of the TIM22 Protein Translocase in Mitochondria. Mol Cell 67, 471–483 e477 (2017).

71. D. Pacheu-Grau et al., Mutations of the mitochondrial carrier translocase channel subunit TIM22 cause early-onset mitochondrial myopathy. Hum Mol Genet 27, 4135–4144 (2018).

72. T. D. Jackson, C. S. Palmer, D. Stojanovski, Mitochondrial diseases caused by dysfunctional mitochondrial protein import. Biochem Soc Trans 46, 1225–1238 (2018).

73. B. Szczesny, T. K. Hazra, J. Papaconstantinou, S. Mitra, I. Boldogh, Age-dependent deficiency in import of mitochondrial DNA glycosylases required for repair of oxidatively damaged bases. Proc Natl Acad Sci U S A 100, 10670–10675 (2003).

74. E. C. Lee et al., A highly efficient Escherichia coli-based chromosome engineering system adapted for recombinogenic targeting and subcloning of BAC DNA. Genomics 73, 56–65 (2001).

75. A. Nagy, M. Gertsenstein, Vintersten, K., R. Behringer, Manipulatiing the Mouse Embryo: A Laboratory Manual. (Cold Spring Harbor Laboratory Press, New York, ed. 3, 2003).

76. M. L. Garcia-Cazarin, N. N. Snider, F. H. Andrade, Mitochondrial isolation from skeletal muscle. J Vis Exp, (2011).

77. V. Dubowitz, R. Lane, C. A. Sewry, Muscle Biopsy: A Pratical Approach. (Saunders Elsevier, Philadelphia, PA, ed. 3rd ed., 2007).

78. R. Patro, G. Duggal, M. I. Love, R. A. Irizarry, C. Kingsford, Salmon provides fast and bias-aware quantification of transcript expression. Nat Methods 14, 417–419 (2017).

79. C. W. Law, Y. Chen, W. Shi, G. K. Smyth, voom: Precision weights unlock linear model analysis tools for RNA-seq read counts. Genome Biol 15, R29 (2014).

80. M. I. Love, W. Huber, S. Anders, Moderated estimation of fold change and dispersion for RNA-seq data with DESeq2. Genome Biol 15, 550 (2014).

81. A. A. Sergushichev, An algorithm for fast preranked gene set enrichment analysis using cumulative statistic calculation. bioRxiv 60012 [Preprint], (2016).

